# A pangenomic perspective of the Lake Malawi cichlid radiation reveals extensive structural variation driven by transposable elements

**DOI:** 10.1101/2024.03.28.587230

**Authors:** Fu Xiang Quah, Miguel Vasconcelos Almeida, Moritz Blumer, Chengwei Ulrika Yuan, Bettina Fischer, Kirsten See, Ben Jackson, Richard Zatha, Bosco Rusuwa, George F. Turner, M. Emília Santos, Hannes Svardal, Martin Hemberg, Richard Durbin, Eric Miska

## Abstract

The East African Rift Lakes, namely Lake Malawi, Victoria, and Tanganyika, host a remarkable diversity of cichlid fishes, representing one of nature’s most striking vertebrate radiations. Despite rich phenotypic diversity, single nucleotide polymorphism (SNP)-based sequencing studies have revealed little sequence divergence between cichlids, with 0.1 to 0.25% pairwise divergence within Lake Malawi. These studies were based on aligning short reads to a single linear reference genome, which ignores the contribution of larger scale structural variants (SVs). To complement existing SNP-based studies, we adopted a pangenomic approach by constructing a multiassembly graph of haplochromine cichlids in Lake Malawi. We produced six new long read genome assemblies, alongside two publicly available ones, to span most of the major eco-morphological clades in the lake. This approach not only identifies longer SVs, but also visually represents complex and nested variation. Strikingly, the SV landscape is dominated by large insertions, many exclusive to individual assemblies. From a pangenomic perspective, we observed an exceptional amount of extra sequence, totaling up to 33.1% additional bases with respect to a single cichlid genome. Approximately 4.73 to 9.86% of the cichlid assemblies were estimated to be interspecies structural variation, suggesting substantial genomic diversity underappreciated in previous SNP-based studies. While coding regions remain highly conserved, our analysis uncovers a significant contribution of SVs from transposable element (TE) insertions, especially DNA, LINE, and LTR transposons. These findings underscore the intricate interplay of evolutionary forces shaping cichlid genome diversity, including both small nucleotide mutations and large TE-derived sequence alterations.

## Introduction

The cichlid fishes in the East African Rift Lakes are one of nature’s most spectacular vertebrate radiations, forming excellent systems to study evolution and speciation (Fryer and Iles 1972; Salzburger 2018; Santos et al. 2023). Lakes Victoria, Malawi and Tanganyika are each home to hundreds of species of cichlids with a wide spectrum of sizes, morphologies, colors, diets and behaviors, shaped by selective pressures in their respective niches (Albertson et al. 2018; Hulsey et al. 2020; Schulte et al. 2014; York et al. 2018). Lake Malawi, in particular, harbors an estimated >800 distinct species predominantly belonging to the Haplochromini tribe (Konings 1989). Cichlids can be categorized into ecomorphological groups based on their genetics (Malinsky et al. 2018). These include the littoral, rock dwelling ‘mbuna’ (1), the generalist *Astatotilapia calliptera* (2), benthics living near the lake floor in shallow (3) or deeper areas (4), the ‘utaka’ species primarily feeding on zooplankton (5), as well as two groups of pelagic cichlids, the deep water *Diplotaxodon* (6) and the piscivorous *Rhamphochromis* (7). Despite the exuberance of phenotypes, single-nucleotide polymorphism (SNP)-based sequencing studies on East African cichlids have revealed extraordinarily low rates of genetic divergence within and between species (Svardal et al. 2020b). Pairwise SNP divergence among Lake Malawi species stood at a mere 0.1-0.25% (Malinsky et al. 2018), approximately ten times lower than that between humans and chimpanzees (Chimpanzee Sequencing and Analysis Consortium 2005).

Because most genomic studies focus on short-read alignments to a single linear reference genome to detect genomic variation, they generally underestimate the contribution of larger scale structural variants, or SVs measuring >50 bp (Mahmoud et al. 2019; Mérot et al. 2020). Growing evidence supports the regulatory roles of SVs in cichlid biology, including in vision (Carleton et al. 2020; Schulte et al. 2014; Nandamuri et al. 2023), sex determination (Munby et al. 2021), body coloring (Kratochwil et al. 2022) and egg spot patterning (Santos et al. 2014), many of which are attributable to transposable element (TE) insertions. For example, syntenic comparisons of the Lake Malawi species *Maylandia zebra* with the much more widely distributed cichlid *Oreochromis niloticus* revealed novel SVs caused by TE insertions (Conte et al. 2019). Another study analyzed SVs across the wider East African radiation (Penso-Dolfin et al. 2020), but utilized highly fragmented Illumina-based assemblies (Brawand et al. 2014), limiting its ability to detect more complex genomic variation. Despite their importance in cichlid adaptive radiations, there is currently limited insight into SVs via a bottom-up, genome-wide approach, especially within individual radiations like Lake Malawi. However, the study of SVs is becoming more accessible with third-generation long-read sequencing technologies such as Pacific Biosciences (Eid et al. 2009) and Oxford Nanopore (Mikheyev and Tin 2014), and we now have high-quality chromosome-level reference genomes for various Lake Malawi cichlid species, including *Astatotilapia calliptera (Rhie et al. 2021)* as well as *Maylandia zebra (Conte et al. 2019)*.

In this study, we adopt a pangenomic perspective to study Lake Malawi cichlid diversity by constructing a multiassembly graph that integrates six newly sequenced assemblies of select haplochromines species, alongside the publicly available genomes of *A. calliptera* and *M. zebra*, spanning most of the major eco-morphological clades. We find the structural variant landscape between cichlids to be dominated by large insertions, many of which are attributable to TEs. We also observe an extensive increase in genomic sequence, totaling up to 33.1% additional bases compared to a single genome when considering the eight genomes together, suggesting substantial previously underappreciated genomic diversity in cichlids.

## Results

### Long read assemblies and graph construction

Six long read assemblies were newly prepared for five Lake Malawi cichlid species, specifically one mbuna (*Tropheops* sp. “mauve”), three benthic (*Aulonocara stuartgranti, Otopharynx argyrosoma*, *Copadichromis chrysonotus)* and one pelagic species (two distinct individuals of *Rhamphochromis* sp. “chilingali”). These genomes were sequenced with either Pacific Biosciences Continuous Long Read (PacBio CLR) technology or Oxford Nanopore (ONT simplex). Further details about sample preparation, sequencing and assembly are provided in the Methods. In addition, we also included the genomes for two other Malawi species downloaded from Ensembl version 103: *A. calliptera* fAstCal1.2, which lives in rivers and marginal areas around Lake Malawi, and another mbuna *M. zebra* M_zebra_UMD2a. Unlike the aforementioned assemblies, these have been scaffolded to chromosome level (scaffold N50 values: *A. calliptera* 38.7 Mbp, *M. zebra* 32.7 Mbp), and they also have gene annotations available. Details about all the assemblies are summarized in Table 1. All eight cichlid genomes, including the publicly downloaded ones, are collapsed, single haplotype assemblies, meaning that we only have information from one of the paternal or maternal alleles at heterozygous sites.

**Table 1:** Lake Malawi haplochromine cichlid genome assemblies. *A. calliptera* and *M. zebra* are chromosome-level assemblies from Ensembl, with values shown in parentheses indicating properties at the scaffold level.

| Species / Sample | Short name | Sex | Sequencing Technology | Assembler | # Contigs / Scaffolds | Contig / Scaffold N50 (Mbp) | Total Size (Gbp) | Sequencing Depth | BUSCO score |
| --- | --- | --- | --- | --- | --- | --- | --- | --- | --- |
| *Astatotilapia calliptera* | astCal | ? | PacBio CLR + Illumina | Falcon | 738 (248) | 4.4 (38.7) | 0.88 | 80.72 | 97.2 |
| *Maylandia zebra* | mayZeb | M | PacBio + Illumina | Falcon | 2331 (1690) | 1.4 (32.7) | 0.96 | 154.12 | 98.4 |
| *Tropheops* sp. "mauve" | troMau | M | PacBio CLR | Falcon | 271 | 8.4 | 0.91 | 137.72 | 98.4 |
| *Aulonocara stuartgranti* | aulStu | M | PacBio CLR | Falcon | 1027 | 2.1 | 0.89 | 86.31 | 98.3 |
| *Otopharynx argyrosoma* | otoArg | M | ONT | Shasta | 2861 | 2.5 | 0.88 | 24.44 | 93.6 |
| *Copadichromis chrysonotus* | copChr | M | ONT | Shasta | 6225 | 0.6 | 0.86 | 19.43 | 92.5 |
| *Rhamphochromis* sp. "chilingali" | rhaChi | M | PacBio CLR | Falcon | 437 | 4.4 | 0.90 | 94.37 | 98.4 |
|  | rhaChi2 | F | ONT | Shasta | 3239 | 1.2 | 0.85 | 42.13 | 93.9 |

The contig N50 values of the newly sequenced genomes range from 0.6 to 8.4 Mbp, with the PacBio assemblies showing better contiguity than their ONT counterparts. Although these genomes are not chromosome-level, they score well when assessed for genome integrity by BUSCO v5.5.0 with the OrthoDB “actinopterygii_odb10” dataset (Manni et al. 2021). Out of the 3,640 essential genes for ray-finned fishes, an average of 3,580 genes (98.4%) were completely matched in the PacBio assemblies, comparable to 3,539 (97.2%) for *A. calliptera* and 3,580 (98.4%) for *M. zebra*, respectively. The ONT assemblies for *O. argyrosoma*, *C. chrysonotus* and *R.* sp. “chilingali” fared worse, with 3,406 (93.6%), 3,366 (92.5%) and 3,420 (93.9%) genes detected, which was expected given that they are more fragmented (**Supplemental Fig. S1**). Nevertheless, these BUSCO scores suggest that our assemblies exhibit overall good integrity when measured by gene completeness. Together, the eight assemblies represent all but one (*Diplotaxodon*) of the six major ecological clades of Lake Malawi cichlids (**Fig. 1A**), which allows a good baseline sampling of the genomic information present in the Lake Malawi radiation.

**Figure 1.**
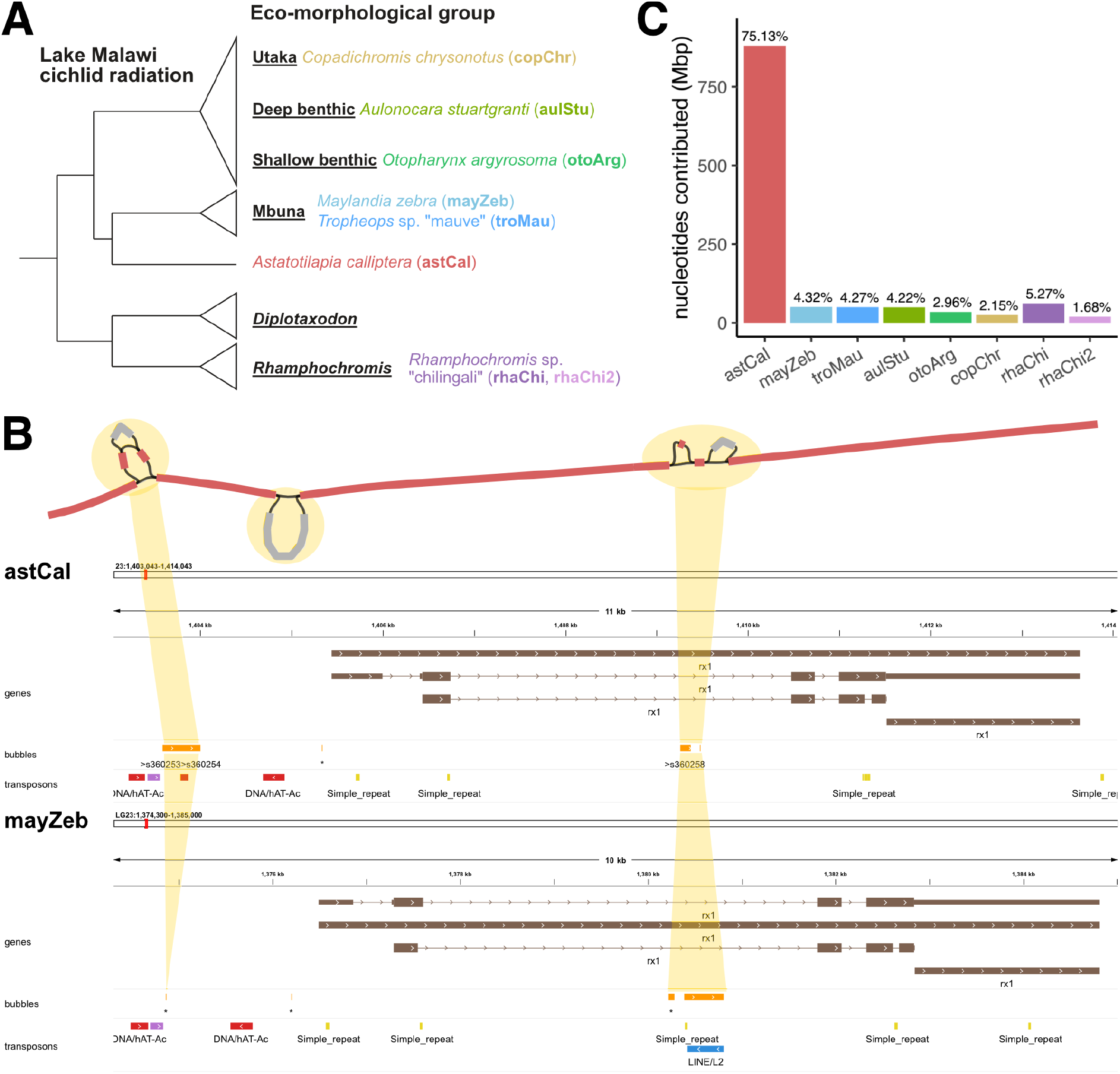
Lake Malawi haplochromine cichlid multiassembly graph. (A) Phylogenetic tree of cichlid species in this study, shown as part of their ecomorphological groups. (B) Graph visualization of the retinal homeobox 1 (rx1) gene, alongside corresponding linear tracks for *Astatotilapia calliptera* and *Maylandia zebra*. (C) Origin of sequence information in graph.

We utilized the minigraph package to construct a multiassembly graph from the Lake Malawi haplochromine cichlid assemblies. Minigraph successively adds new genome sequences to the graph structure by genome-to-graph alignment, starting from a reference genome assembly, known as the backbone (Li et al. 2020). Bubbles are augmented onto the existing graph to represent regions where the new assembly sufficiently differs in sequence from all those already incorporated. We chose the *A. calliptera* (astCal) genome as the backbone because it is the most contiguous assembly with Ensembl gene annotations available, allowing us to define where SVs are positioned relative to known genes. The remaining assemblies were incorporated in the following order, prioritizing phylogenetic proximity and genome quality: *M. zebra* (mayZeb), *T.* sp. “mauve” (troMau), *A. stuartgranti* (aulStu), *O. argyrosoma* (otoArg), *C. chrysonotus* (copChr) and the two *R.* sp. “chilingali” (rhaChi, rhaChi2).

### Structure of the Lake Malawi haplochromine cichlid multiassembly graph

The Lake Malawi cichlid multiassembly graph consisted of linear structures representing the backbone scaffolds, which are punctuated by bubbles representing SVs (**Fig. 1B**). Overall, the graph contains 1,171,902,121 nucleotides, which were distributed across 637,237 segments connected by 913,087 edges. On average, a segment measured 1,839.04 nucleotides in length and was connected by 1.43 edges. The linear part of the graph comprised 189,197 segments containing 758,854,960 bases (64.8% of the total), while the variable part (bubbles) was made of 448,040 segments containing 413,047,161 bases (35.2%). As the query assemblies were incorporated, the graph showed greater complexity with increases in the number of edges and segments, accompanied by a decrease in the average length of segments (**Supplemental Fig. S2**). The backbone-guided nature of graph construction in minigraph meant that the majority of the graph segments originated from the *A. calliptera* backbone, whose 880,428,986 nucleotides were encompassed within 382,935 segments. Incrementally incorporating the assemblies for mayZeb, troMau, aulStu, otoArg, copChr, rhaChi and rhaChi2 grew the reference graph by 62,307, 40,627, 44,206, 29,209, 21,444, 40,742 and 15,767 segments, which consisted of 50,651,908, 50,003,806, 49,458,672, 34,661,015, 25,161,891, 61,804,019 and 19,731,824 nucleotides respectively (**Fig. 1C**). The query assemblies contributed a cumulative total of 291,473,135, or 33.1%, additional nucleotides compared to the *A. calliptera* backbone. These nonreference sequences were encapsulated within a total of 188,944 bubbles, positioned in backbone regions that span the structural breakpoints from at least one of the nonreference samples. Out of the 880,428,986 nucleotides on the backbone segments, a substantial 840,567,200 (95.47%) received spanning coverage (**Supplemental Fig. S3, S4**), which translates to a bubble density of 0.2248 per 1 kbp of covered backbone sequence.

The construction of a multiassembly graph with minigraph relies on the initial choice of a backbone assembly, and it has been reported that this could directly impact the amount of nonreference sequence that is detected from the subsequent assemblies (Crysnanto et al. 2021). To investigate the robustness of our multiassembly graph, we built seven other graphs using each of the nonreference samples as the backbone in turn. The more contiguous PacBio backbones produced graphs that were topologically more complex than their ONT counterparts with larger number of segments, edges, bubbles and extra sequence. Nevertheless, the proportions of linear and variable sequences were similar across all the backbones at about approximately 65% and 35% respectively (**Supplemental Fig. S5**), while most metrics in the PacBio-based graphs fell within 5% of those of the canonical *A. calliptera* version (**Supplemental Table S1**). Permuting the order of incorporation of the subsequent assemblies did not significantly affect graph structure and topology, suggesting that backbone choice was the dominant factor in the variation that was observed (**Supplemental Fig. S6**). For a fair comparison of bubble frequency, we normalized the bubble counts by the number of backbone nucleotides with spanning coverage, and found that these values ranged from 0.2192 to 0.2254 bubbles per kbp of covered genomic sequence (*A. calliptera*: 0.2248). These observations suggest that minigraph is consistent for structural variant discovery, and that the default graph exhibited a reasonable structure and amount of sequence information.

Finally, we examined how multiassembly graph construction behaved in response to the choice of the minimum variant length (L) in minigraph, which influences the minimum length of sequence difference required for a bubble to be created. The default value of L = 50 follows the conventional definition of a structural variant (SV), which ignores smaller scale variation (SNPs, small indels) and makes the graph less complex and more interpretable. We think that the default is appropriate for our biological hypothesis and the quality of our assemblies. Although the topological complexity of the multiassembly graph decreases with larger values of L, we found that there was a relatively broad parameter range of L = 25 to 500 where the total sequence content in the reference graph remained relatively stable (**Supplemental Fig. S7, Supplemental Table S2**).

### Rapid growth of nonreference sequences

An interesting property of our Lake Malawi cichlid multiassembly graph is the increasing growth in the amount of variable sequences with each incorporated nonreference assembly, displaying a sharp incline with indications of converging towards an asymptote (**Fig. 2A**). After all the samples were integrated into the graph, the amount of nonreference sequence contributed by the nonreference assemblies totalled up to 33.1% the size of the *A. calliptera* backbone. Conversely, the flexible sequence on the *A. calliptera* backbone already showed signs of plateauing at around 13.81% (121.57 Mbp). Repeating this analysis across the graphs with different Malawi backbones also showed this rapid growth in flexible sequences attributed to the nonreference assemblies, rather than more of any backbone being uncovered as such (**Supplemental Fig. S8**). This suggests that each assembly contains additional stretches of genomic sequences that might have been overlooked if we had only focused on any individual backbone as a reference.

**Figure 2.**
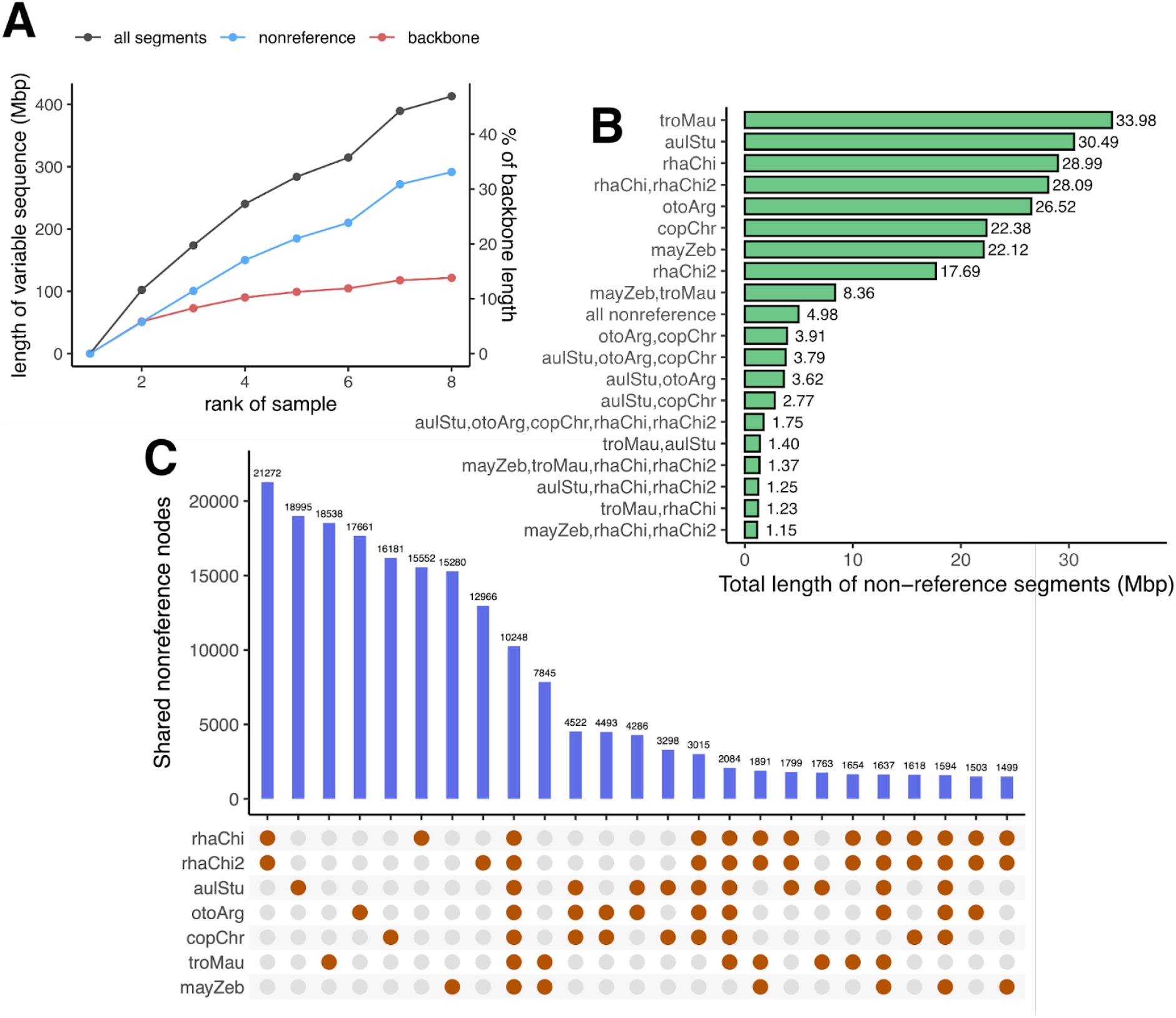
Nonreference sequences in Lake Malawi cichlids. (A) Growth of the variable components in the graph, based on whether they originated from nonreference assemblies (blue) or the backbone (red). (B, C) Cumulative length and number of segments shared across assemblies.

Consistent with this, we uncovered that a majority of nonreference segments in the bubbles are private to one assembly, also known as singletons (**Fig. 2B, 2C**). Correspondingly, there was not a single assembly from which a significant majority of the nonreference segments could be traced, with each assembly contributing a similar share. The three largest contributors were the higher quality PacBio assemblies of troMau (33.98 Mbp), aulStu (30.49 Mbp) and rhaChi (28.99 Mbp) from the mbuna, benthic and *Rhamphochromis* groups respectively. These singletons cumulatively account for 182,158,018 (62.5%) out of 291,473,128 nucleotides spread across 115,173 (45.3%) out of the 254,201 nonreference segments present within the bubbles. The greatest amount of sequence shared between at least two nonreference assemblies came from the two *Rhamphochromis* assemblies with 28.09 Mbp shared across 21,272 segments. Sharing between all samples of the same ecological clade was not as extensive, with the mbuna (mayZeb and troMau) sharing 8.36 Mbp across 7,845 segments, and the benthic group sharing 3.79 Mbp across 4,522 segments. There was 4.98 Mbp of sequence located on 10,248 segments which were missing from *A. calliptera* but shared by all the nonreference assemblies. These bubbles conversely contained a substantial amount of backbone sequences which were private to the *A. calliptera* assembly: 22,511 segments harbouring 42.32 Mbp (4.81% of the backbone). These results suggest that the structural variant landscape in our Lake Malawi cichlid multiassembly graph is not only dominated by singletons contributed by augmenting the backbone with nonreference assemblies, but there are also unique sequences present in the backbone that we identified through the pangenomic approach.

### Discovery of structural variants

We investigated the 187,552 bubbles in the reference graph in detail, and found that the majority harbored straightforward present-or-absent variation or slightly nested variation, with complex structural variation being relatively rare (**Supplemental Fig. S9**). The number of possible paths to transverse a bubble exhibited a strongly left skewed distribution towards simple variation: there were 151,761 (81%) 2-path and 21,230 (11%) 3-path bubbles, and 182,136 (97%) had no more than eight theoretical paths (**Supplemental Fig. S10**). To focus on true biological variation within the bubbles, or the alleles, we retain only the paths actually transversed by samples. Using minigraph, we were able to successfully call alleles for all samples at 160,578 out of the 188,452 bubbles (85.2%). For subsequent analyses, we retained the 187,552 out of 188,452 bubbles (99.5%) for which the alleles are known for at least five out of eight samples to facilitate between-sample comparisons (**Supplemental Fig. S11**).

Most of the retained bubbles were bi-allelic (153,848, 82%), while the remaining multi-allelic ones mostly contained three alleles (20,751, 11%). The graph was composed of mostly simple structural variation, many of which were singletons (**Fig. 3A**), but the size of the SVs varied widely. The lengths of structural variant alleles exhibited a bimodal distribution, with one peak at around 50 bp and another above 1,000 bp. A total of 434,357 alleles were observed across the 187,552 bubbles, which translated to an average of 2.32 alleles per bubble. The 246,805 nonreference alleles in this set comprised 144,769 insertions, 101,179 deletions and 857 substitutions/inversions with an average length of 2016.7, 1510.8 and 605.2 bases, respectively (Fig. 3B). The cumulative length of insertions was 291,957,550 bases, which was longer than the 152,861,637 associated with deletions. The Lake Malawi cichlid multiassembly graph contained more complete insertions (78,479; only nonreference allele with sequence, reference length of 0) compared to partial/alternate insertions (66,290; both reference and nonreference sequences present, but the nonreference allele is longer). We observed the opposite pattern with deletions: 49,217 complete vs 51,962 partial/alternate.

**Figure 3.**
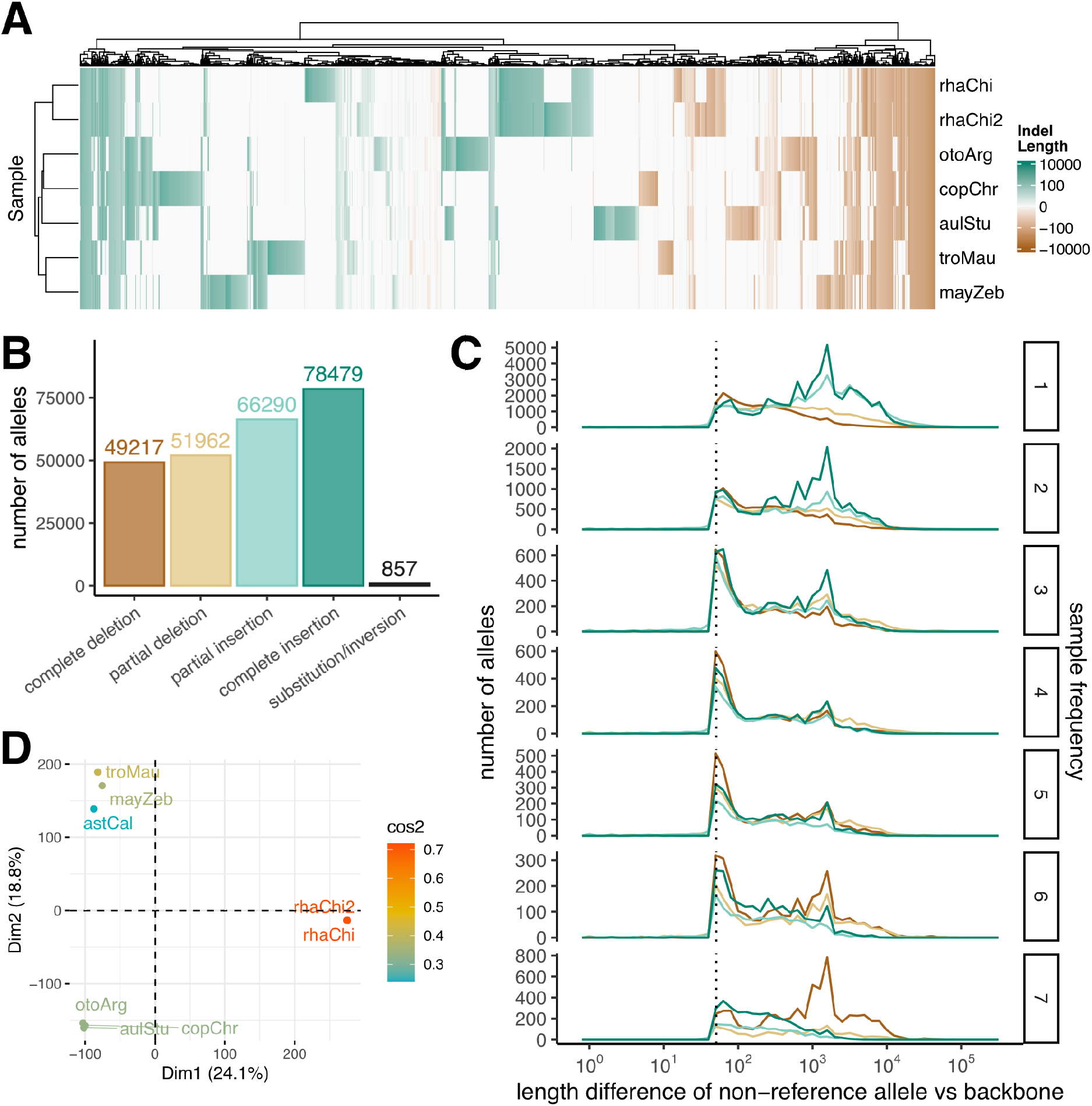
The Lake Malawi cichlid structural variant landscape. (A) Heatmap showing length difference of alleles for each nonreference sample versus *A. calliptera* backbone across 5000 random bubbles. A positive value (green) represents an insertion, while a negative value (brown) represents a deletion, with darker colors indicating larger SVs/indels. (B) Number of structural variant alleles by type. (C) Length deviations of nonreference alleles across different sample frequencies. Dashed line denotes 50 bp. (D) PCA performed with allele lengths at the 160,572 bubbles with complete allelic information.

Most of the biological alleles in the reference graph are single insertion or deletion events that are present at a low sample frequency across our eight assemblies. A significant majority of alleles (147,802 or 60%) are singletons, meaning that the nonreference allele either had a sample frequency of 1 out of 8 (135,160 alleles), or 7 out of 8 (12,642 alleles), in which case the reference allele on the *A. calliptera* backbone is the singleton (**Fig. 3C**). These results are consistent with the observation in the previous section that many nonreference segments and sequences are private to a single assembly (**Fig. 2**). The extreme sample frequencies were primarily dominated by the large structural variants measuring thousands base pairs in length, with 1/8 and 2/8 frequencies dominated by complete insertions, while 7/8 by complete deletions (i.e. complete insertions in *A. calliptera* backbone relative to the others). Conversely, intermediate allele frequencies tended to be composed of shorter sequence variants, with most having a length of around 50 bp (**Fig. 3C**).

### Recapitulation of species relationships

Next, we investigated how the phylogenetic information in the multiassembly graph compared to what was known about species relationships from SNP studies of the Lake Malawi cichlid radiation (Malinsky et al. 2018). Using the biological allele lengths in the 160,572 bubbles with complete allelic information, we successfully separated the samples by ecological clade in a principal component analysis (PCA) plot (**Fig. 3D**), and they showed the expected clustering based on Pearson correlation values and on the number of sites with different alleles (**Supplemental Fig. S12**). The two *Rhamphochromis* individuals showed closer clustering than any other pair of assemblies, scoring a Pearson’s correlation of 0.74, compared to approx. 0.51 within benthic and 0.59 between mbuna samples. Moreover, the *Rhamphochromis* assemblies differed in allele sequences at 31,180 sites which was half of those in other pairs. Overall, this suggests that our multiassembly graph was phylogenetically sound and successfully recapitulated the expected interspecies relationships based on the prevailing understanding of the evolutionary history of the Lake Malawi cichlid radiation, despite the fact that the assemblies vary in their quality and the sequencing technologies used.

As a further analysis on the cichlid relationships, we utilized minigraph to build bi-assembly graphs from every possible pair of species, allowing a fair comparison of any pair of assemblies, without any potential alignment bias caused by the augmentation of bubble sequences from the other assemblies. This comparison is asymmetrical, where one assembly acts as a backbone and the other as a query, from which an estimate of the sequence divergence caused by SVs is obtained by calculating the backbone regions with spanning coverage which were encapsulated inside bubbles (**Fig. 4A**). With the default minimum variant length parameter L = 50, estimated interspecies SV divergences ranged from 4.73% to 9.86% (mean: 7.11%), with smaller values within the mbuna (mean: 5.45%) and benthic (mean: 5.20%) clades. The estimated divergence values were much lower within species at 2.96% and 3.68% when comparing the two *Rhamphochromis* individuals. We also tested parameter values of L from 2 to 250, along which smaller variants were gradually excluded, causing the percentage conservation scores to decrease and eventually reach a plateau (**Supplemental Fig. S13**). The genome graphs generated at higher L values should be more reliable as they exclude small sequencing errors due to the relatively high error rates in PacBio CLR and ONT simplex reads that were used for assembly.

**Fig. 4:**
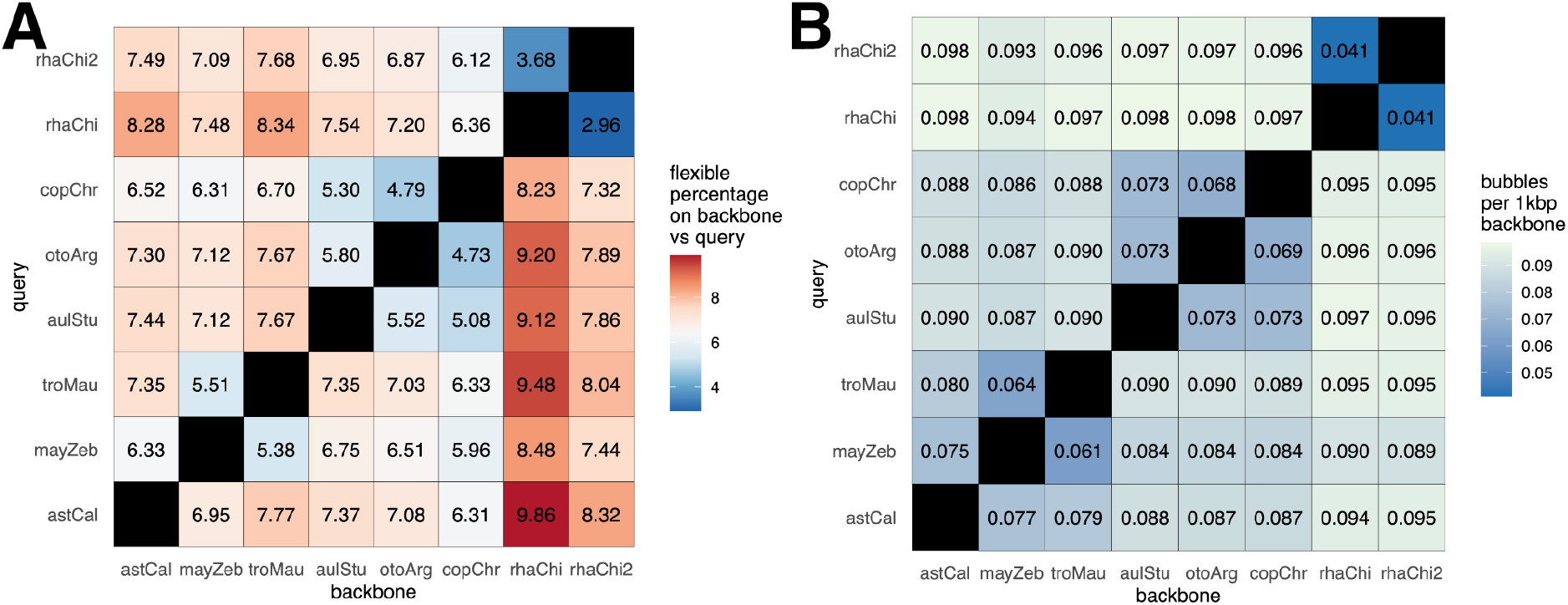
Estimated pairwise SV sequence divergence (a) and bubble density (b) between Lake Malawi cichlid assemblies. Values were calculated using bi-assembly graphs between pairs of cichlid assemblies. Minimum variant size in minigraph was set to 50.

Using these same bi-assembly graphs, we next obtained a measure of how frequently structural events occur along the genome by calculating the bubble density normalized by spanning coverage (**Fig. 4B**). This revealed an interspecies bubble density of 0.061 to 0.098 per 1 kbp of backbone, compared to an intraspecies density of 0.041 among the two *Rhamphochromis*. Interestingly, there appeared to be some directionality to the sequence changes, as both *Rhamphochromis* individuals contained more insertions and fewer deletions when aligned to the other species (**Supplemental Fig. S14**). This high similarity between the two *Rhamphochromis* individuals is expected since they belong to the same species and are closely related (both individuals originate from the same aquarium stock). The *Rhamphochromis* assemblies exhibit structural changes independently of the other cichlids, which is consistent with their positioning in the SNP-based Lake Malawi cichlid phylogeny, where the pelagic clade that contains the *Rhamphochromis* and *Diplotaxodon* taxa branched off earlier than the rest of the radiation.

### PCR validation and genotyping

We performed PCR (polymerase chain reaction) validation of 16 predicted SVs, using five of the original tissue samples that were used for genome sequencing of: troMau, otoArg, copChr, rhaChi and rhaChi2. Because of difficulties in designing PCR primers for repetitive genomic regions, we prioritized bubbles containing simple, straightforward presence-or-absence variants located within 1500 bp of a gene. Out of the sixteen bubbles we tested, the validation results for twelve were consistent with the allele predictions for all five assemblies, while the remaining four scored correct for four assemblies (**Fig. 5A, Supplemental Table S3**). Some heterozygosity was present in four of the twelve bubbles, which was expected because our samples were not from homozygous lines and therefore will have contained heterozygous SVs where the two alleles differ, whereas their single haplotype assemblies only contained one of these two alleles. 10 out of the 16 SV PCR results were further verified by Sanger sequencing (**Supplemental Table S3**). For the remaining 6/16 SVs, PCR results were partially validated by Sanger sequencing, as we could not acquire good quality sequencing in some instances, likely due the repetitive nature of these loci. However, all sequenced PCR products that were obtained corresponded to the expected sequences. Overall, these results provided additional experimental support to our finding, and demonstrate that minigraph’s pangenomic approach to detecting structural variants was reliable.

**Figure 5.**
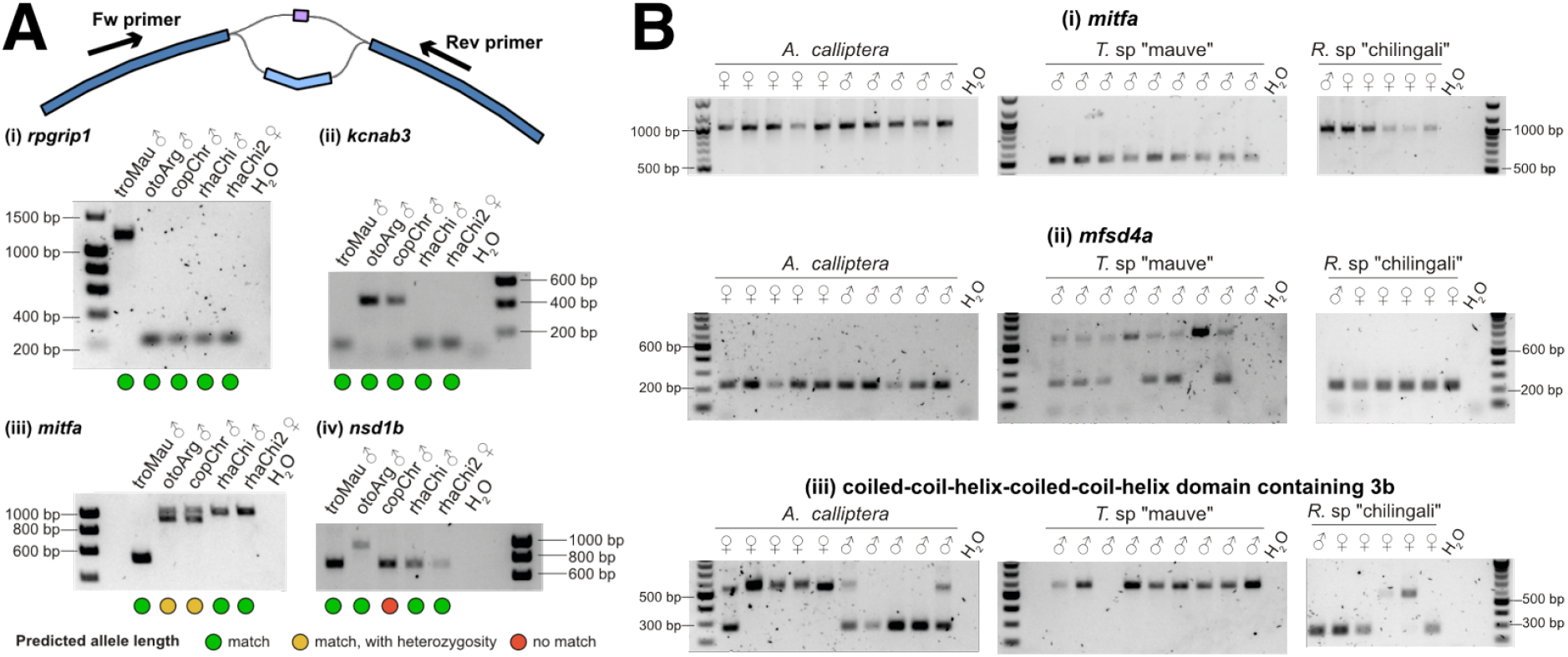
PCR genotyping of structural variants. (A) Validation of predicted SVs in original samples. Green = match; amber = match with heterozygosity; red = no match. (B) Wider genotyping across aquaria-grown individuals: 10 *A. calliptera* (5 males, 5 females), 9 *Tropheops* males and 6 *Rhamphochromis* (1 male, 5 females).

We also utilized the designed PCR primers to perform additional genotyping of a wider cohort of aquaria reared male and female individuals of *A. calliptera*, *T.* sp. “mauve” and *R.* sp. “chilingali” to check for evidence of polymorphism at some bubbles (**Fig. 5B**). For example, we did not find any evidence for polymorphism in a bubble near *mitfa* (ENSACLG00000022427) with all genotyped *A. calliptera* and *R.* sp. “chilingali” showing one allele, and all *T.* sp. “mauve” showing the other. However, bubbles near some genes (*mfsd4a* and ENSACLG00000002767) showed evidence of polymorphism in different species. These observations suggest a variable degree of polymorphism and heterozygosity for structural variants across the different sexes and species. However, there is limited statistical power to draw any conclusions about the wider Lake Malawi cichlid population given our small sample sizes.

### Genes

To investigate the functional relevance of the structural variants, we examined the 187,552 bubbles for the presence of nearby genes. We focused on a subset of 26,734 out of 28,001 gene annotations in the *Astatotilapia calliptera* genome, filtering out putative TEs misannotated as genes (see Methods). 105,508 (56.3%) bubbles were either located within or in close proximity to a gene: 90,531 (48.3%) directly intersected with genic features like exons, introns or UTRs; while 8,967 (4.8%) and 6,010 (3.2%) bubbles were located within 2,000 bp upstream and downstream of genes, respectively. The remaining 82,024 (43.7%) were considered intergenic, with large average distances from the nearest gene start and end at 38.4 and 41.2 kbp respectively. Coverage-corrected bubble densities were 0.2481 per kbp of intergenic sequence and 0.2213 for introns, similar to the genome-wide level of 0.2248, but the bubble density in exonic regions was significantly lower at 0.0899 per kbp (**Fig. 6A**). 10,364 (38.8%) out of 26,734 genes were devoid of any SVs in the gene body, and did not contain any bubbles in the regions 2,000 bp upstream (57.9%) and downstream (69.5%). Genes containing SVs are associated with transport (*p* = 2. 122 × 1 0^−33^), phosphorus metabolic process (*p* = 6. 204 × 1 0^−23^) and developmental process (*p* = 5. 660 × 10) (**Supplemental Fig. S15**). There were 4,723 (17.7%) genes where SVs were entirely absent from the gene body and within a 2,000 bp region, which we consider a highly conserved set of genes across species. This gene set was associated with Gene Ontology terms pertaining to processes in gene expression, transcription and translation: gene expression (*p* = 4. 4 × 1 0^−40^), RISC complex (RNA-induced silencing complex, *p* = 9. 2 × 10), DNA-binding transcription factor activity (*p* = 1. 6 × 10) and structural component of ribosome (*p* = 1. 9 × 1 0^−10^) (**Supplemental Fig. S16**).

**Figure 6.**
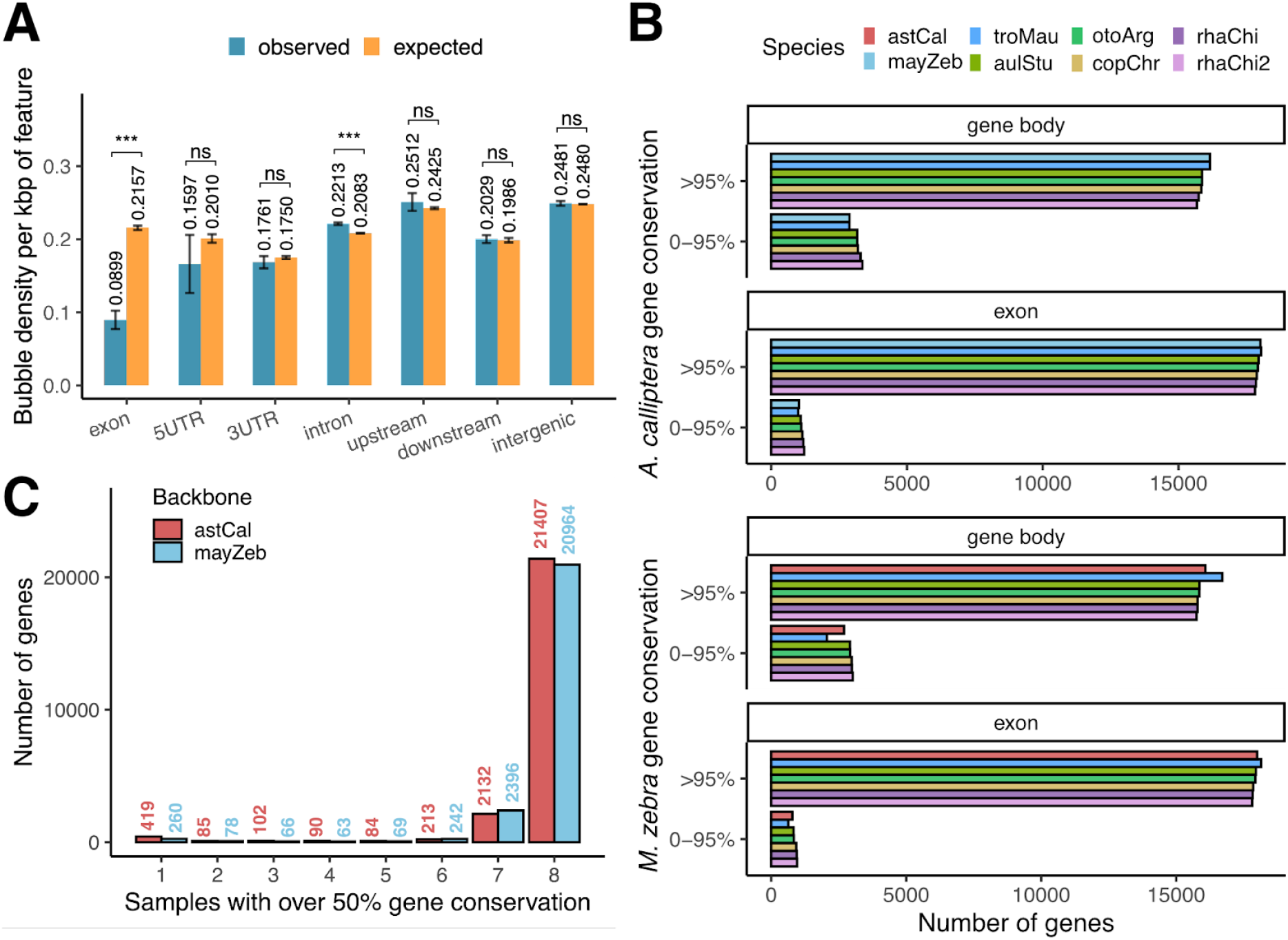
Genomic context of structural variants. (A) Bubble density in gene features. 95% confidence intervals are estimated by bootstrapping. *** *p* < 1 × 10 ^−3^, n.s. not significant. (B) Percentage sequence conservation of genes calculated for backbones with gene annotations. (C) Presence-absence counts of backbone genes. A value of 1 denotes genes not detectable in the other assemblies and might be private, while a value of 8 refers to ubiquitous genes.

Subsequently, we implemented a graph-oriented approach with the ODGI package to estimate the percentage sequence of each gene present in each sample. This was achieved by overlaying gene coordinates onto graph segments, and calculating a conservation score based on the path taken by its assembly through the graph. We performed this presence-or-absence variation (PAV) analysis for the chromosome-level *A. calliptera* and *M. zebra*-backboned graphs, which had gene annotations available, to account for possible reference bias because the backbones will score almost perfect conservation of genes on their own graphs. We also restricted the analysis to a set of 24,322 (86.9%) and 24,532 (85.6%) genes in *A. calliptera* and *M. zebra*, respectively, falling in genomic regions without overly complex bubbles and spanning coverage from at least six samples, and removed potential transposon misannotations. Through this approach, we discovered a high degree of conservation in protein-coding genes. Specifically, 89.1 to 90.8% of *A. calliptera* genes demonstrated at least 95% sequence conservation in the gene body across the samples, with ecologically similar species exhibiting similar percentage conservation scores to each other (**Fig. 6B**). This conservation was more pronounced in exonic regions, where 94.9 to 95.5% of genes maintained this high level of sequence similarity (**Supplemental Table S4**).

These sequence conservation scores also allowed us to tally how frequently each gene was present across the eight assemblies, where “present” denoted a gene with 50% of its sequence being detectable in a given sample. This threshold was fairly lenient to account for the fact that genes can have very large introns and 4.73-9.86% of genomic regions could manifest as SVs between species (**Fig. 4A**). As anticipated, a large proportion of gene sequences from 21,407 (88.0%) *A. calliptera* and 20,964 (85.5%) *M. zebra* were detectable in the other assemblies. There was substantial concordance between these gene sets based on orthology information from BioMart: 19,850 (93.2%) of *A. calliptera* genes had corresponding homologs in *M. zebra*, and 19,626 (93.6%) vice versa. This PAV approach also suggested that *A. calliptera* and *M. zebra* respectively harbor 378 and 260 gene sequences not detectable in the other assemblies (**Fig. 6C**). However, by cross-referencing orthology data from BioMart, more than half of these apparently private genes were found to actually contain homologs in the other assembly, leaving only 150 *A. calliptera* and 94 *M. zebra* gene sequences that might be authentic “backbone-specific” sequences. Gene Ontology functional enrichment analysis of these were not informative, but a Pfam domain alignment search (**Supplemental Fig. S17**) indicated the presence of protein domains that are fairly ubiquitous across eukaryotic protein families, including ubiquitin-like (e.g. ubiquitin, Crinkler), zinc-fingers, ankyrin repeats, thrombospondin type-1 repeat (TSP1), trypsin inhibitor-like cysteine-rich (TIL) and G-protein-coupled receptor (GPCR) domains. Transposon-associated domains such as endonuclease, reverse transcriptase and transposase were also detected.

The sets of genes in *A. calliptera* (228) and *M. zebra* (166) that were PAVs in the graph but for which Biomart orthologs are found could indicate genomic regions where there might have been assembly errors or technical artifacts that were misinterpreted as structural variation during graph construction (**Supplemental Fig. S18**). Although transposon-related domains were present, we also detected the presence of additional protein-coding domains in these sets, the most common of which belong to immunoglobulins (e.g. V-set, I-set and Ig domains). Noteworthy was the appearance of several hemoglobin genes supposedly private to the *A. calliptera* assembly, accompanied by the Gene Ontology terms hemoglobin complex (*p* = 2. 7 × 1 0^−3^) and oxygen carrier activity (*p* = 3.3 × 1 0 ^−2^). However, inspection in IGV suggests that these might have been caused by assembly errors around the hemoglobin MN locus on the *A. calliptera* backbone, causing the formation of artifact bubbles when the other cichlid assemblies were aligned (**Supplemental Fig. S19**). The inherently repetitive nature of sequences within TEs, immunoglobulins, and hemoglobins poses challenges for accurate genome assembly, and these false positives emphasize the need for better assemblies and annotations for more robust conclusions. Nevertheless, our available evidence suggests that the structural divergence of Lake Malawi cichlids is unlikely to have been caused by preferential gene gain or gene loss in certain clades.

### Transposable elements

Having found that genes are not strongly related to the SVs between the assemblies, we turned our attention to TEs, which have been shown to be sources of phenotypic novelty in cichlid fishes (Santos et al. 2014; Carleton et al. 2020; Munby et al. 2021). About 36.55% of the *A. calliptera* sequences are annotated by RepeatModeler/RepeatMasker as TEs, of which the biggest classes were DNA transposons (12.09%), LINEs (8.32%) and LTR retrotransposons (7.38%), while the SINEs, Helitrons and Retroposons collectively take up less than 1% (**Figure 7A**). There is also a substantial amount of Unknown elements (8.54%), which might constitute currently uncharacterised transposons and other repetitive elements. Overlap analysis suggested that there was noticeable enrichment of TEs in structural variant regions on the backbone at a percentage sequence of 74.65%, a 2.04-fold enrichment compared to genome wide, or 2.20-fold if Unknown elements were excluded (62.25% TEs). This enrichment was detected when SV coordinates were randomized and across varying sequence divergence thresholds for TE detection in RepeatMasker, with the DNA, LINE and LTR transposons consistently showing strong enrichment (**Supplemental Figs. S20, Supplemental Table S5**). These genomewide and SV transposon proportions were also reflected in the other species: ranging from 36.32 to 38.29% genomewide, and 72.26 to 75.74% in SVs, suggesting that TEs represent most of the large scale sequence differences among the cichlid assemblies (**Supplemental Table S6**).

**Figure 7.**
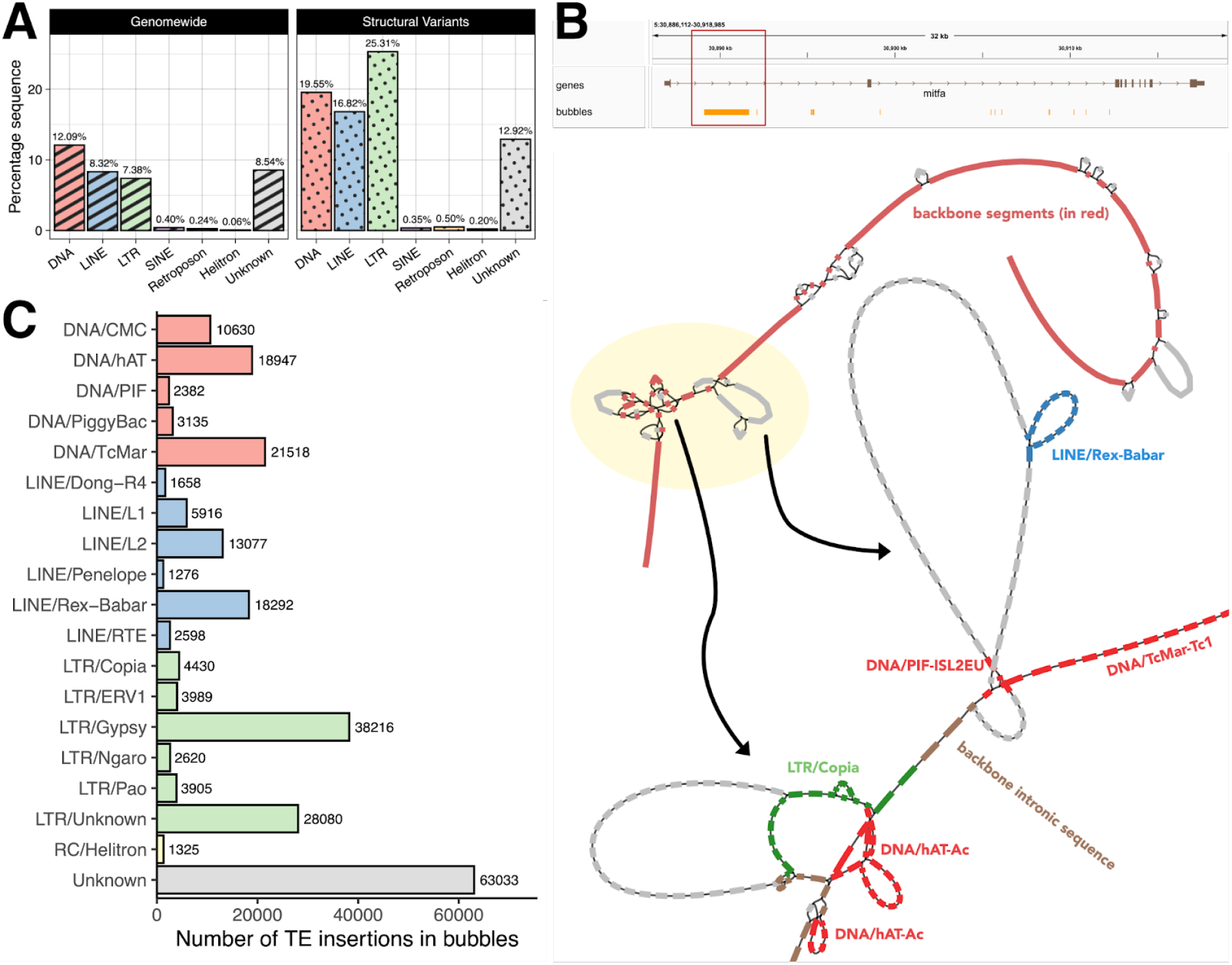
Genomewide and structural variant transposon composition of the *Astatotilapia calliptera* fAstCal1.2 assembly. (B) TE insertions in the first intron of *mitfa*, highlighting the presence of two complex bubbles. (C) Number of putative polymorphic TE insertion events by subclass across 90,451 bubbles.

Closer inspection showed that the multiassembly graph successfully incorporated known examples of SV/TEs in Lake Malawi cichlids, such as a haplochromine-specific SINE insertion upstream of *fhl2b* (**Supplemental Fig. S21**) (Santos et al. 2014) and a nested insertion upstream of *rx1* whose alleles contribute to different opsin palettes in cichlid vision (**Supplemental Fig. S22**) (Schulte et al. 2014). Another example is the *mitfa* gene, whose first intron harbors the presence of complex variation caused by TE insertions, consistent with previous observations (Carleton et al. 2020). While we refrain from making claims about species-level differences in TE composition, due to the varying quality of the assemblies, we performed a high-level characterization of the subclasses of TEs within graph bubbles as they might indicate classes that are still segregating within the Lake Malawi population. By overlapping TE coordinates against SVs across all the assemblies, we identified 254,015 straightforward present-or-absent TE insertion events across 90,451 unique bubbles (**Fig. 7C**). The most frequent within-bubble TE insertions events involved the subclasses of LTR/Gypsy (38,216), LTR/Unknown (28,080), DNA/TcMar (21,518), DNA/hAT (18,947) LINE/Rex-Babar (18,292) and LINE/L2 (13,077). On a genome-wide scale, 14,868 genes (55.9%) had a putative polymorphic TE bubble in the gene body or within 2,000 bp upstream. This number decreased to 7,898 (29.7%) if we specified an additional constraint requiring the bubble to be in proximity to the start of the gene. None of these intersections revealed a significant Gene Ontology enrichment, even when stratifying for TE classes and subclasses. This lack of functional enrichment supports the hypothesis that species-variable TE insertions display no apparent pattern or preference for specific genes.

## Discussion

By integrating eight long-read genome assemblies of Lake Malawi haplochromine cichlids into a multiassembly graph, we uncover and characterize novel structural variation not represented in the established chromosome-level assemblies of *A. calliptera* and *M. zebra*. We estimate that there is 26.4 to 33.1% of additional sequence relative to a chosen assembly (Fig. 2), a relatively large amount compared with reports from intraspecific reference graphs, including cattle (2.8% in 6 genomes (Crysnanto et al. 2021)), sheep (∼5% in 15 breeds (Li et al. 2023)), duck (2.33% flexible genes in 131 genomes (Wang et al. 2023)) and humans (∼10% in 910 African individuals (Sherman et al. 2019)). Direct comparisons of these values are not straightforward due to variations in methodologies and lack of universal definitions of pangenomic terms (Sherman and Salzberg 2020), but this study supports the existence of a significant amount of underappreciated sequence diversity in cichlids. Although structural variants have been studied before in cichlids (Brawand et al. 2014; Conte et al. 2019; Fan and Meyer 2014; Kratochwil et al. 2019; Penso-Dolfin et al. 2020), as far as we are aware, our work not only represents the first efforts to quantify SVs in these fishes using a pangenomic approach, but also elucidates these large-scale genomic differences within a single haplochromine tribe of cichlids in a single East African lake, which we hope will pave the way towards more comprehensive pangenomic references for cichlids. We estimate that the interspecies pairwise divergence between our assemblies range from 4.73 to 9.86% of base pairs in SVs (Fig. 4), contrasting with the 0.1 to 0.25% values quoted for SNPs (Malinsky et al. 2018). These numbers represent different aspects of genomic variation. SNP divergence provides not only a direct measure of the percentage sequence that differs, but also indicates the number of events causing these differences (e.g. 0.1% denotes on average one event per 1000 bp of sequence). In contrast, our calculated SV divergence values specifically estimate the percentage sequence of SVs in a given genome, but the number of events is often smaller. These metrics unveil distinct aspects of cichlid genome biology: SNPs are useful for assessing genetic relatedness (Malinsky et al. 2018), performing genome-wide association studies (Kratochwil et al. 2022) and scanning for signatures of selection (Malinsky et al. 2015), while our SV analysis was useful for variation in intronic and intergenic regions outside protein-coding genes. It is not surprising that most SVs are relatively less common within coding segments of genes, as such alterations within coding regions can be highly detrimental, potentially disrupting the reading frame or introducing premature stop codons (Almeida et al. 2022). Despite these differences, it is worth emphasizing that both SVs and SNPs likely exhibit similar population-level patterns in cichlid radiations, with evidence of polymorphism within and between species, as shown for select SV alleles in our PCR genotyping experiments (Fig. 5). This implies that although we have identified certain non-reference sequences as occurring only once in an assembly, future studies will need to obtain population level SV information to assess within-population SV allele frequency distributions.

Although the common ancestor for East African haplochromine cichlids has been inferred to have a high gene duplication rate relative to other teleosts (Brawand et al. 2014), our cichlid assemblies do not allow us to confidently conclude whether the descendant ecomorphological clades within Lake Malawi show preferential gains or losses in eukaryotic gene families. Minigraph’s whole-genome alignment approach produces a more straightforward graph to visualize complex variation (Li et al. 2020), but it requires that the incorporated genome assemblies are reliable and accurate to begin with. Errors within the backbone assembly might encapsulate backbone gene annotations within artifactual bubbles, falsely suggesting the absence of certain genes in other assemblies. An example was illustrated by an assembly error resulting in a technical duplication of a hemoglobin locus in the *Astatotilapia calliptera* assembly (**Supplemental Fig. S19**). We anticipate a decrease in such problems as newer assemblies with improved accuracy become available (Rhie et al. 2021).

The pangenomic graph approach emerges as a computationally efficient method for comparing multiple whole genome assemblies when characterizing TE insertions at sites with complex variations (Ebler et al. 2020; Groza et al. 2023). Notably, our findings reveal a striking 74.65% of structural variant regions attributable to TEs. While prior research has uncovered isolated instances of beneficial or co-opted TE insertions in cichlids (Santos et al. 2014; Carleton et al. 2020; Munby et al. 2021), our results suggest a widespread occurrence of such insertions in the Lake Malawi cichlid genomes. Although the majority of TE insertions are expected to be evolutionarily neutral (Arkhipova 2018), it is intriguing to speculate that persistent TE activity might serve as a mechanism for closely related species like cichlids to maintain sufficient genomic differences, giving potential to produce diverse phenotypes if coincided with ecological opportunity (Ngoepe et al. 2023). This may occur even in the absence of SNP differences resulting from pervasive gene flow (Svardal et al. 2020a; Meier et al. 2023), although future studies should attempt a population genetics approach to ascertain if similar dynamics are at play. Nevertheless, our findings underscore the complex interplay of evolutionary forces shaping the Malawi cichlid genome, where diversity is the product of not only SNPs and small-scale differences caused by point mutation, but also structural variation driven primarily by TE activity.

## Materials and Methods

### Genome assemblies

A total of eight Lake Malawi cichlid long read genomes were used: two previously published chromosome-level assemblies from Ensembl v103 — *Astatotilapia calliptera* (fAstCal1.2, GCF_900246225.1) (Rhie et al. 2021) and *Maylandia zebra* (M_zebra_UMD2a, GCA_000238955.4) (Conte et al. 2019), as well as six contig-level assemblies generated for five other species from aquaria-grown fishes — PacBio CLR for *Tropheops* sp. “mauve”, *Aulonocara stuartgranti* (male), and *Rhamphochromis* sp. “chilingali” (male); and ONT simplex for *Otopharynx argyrosoma* (male), *Copadichromis chrysonotus* (male), and *Rhamphochromis* sp. “chilingali” (female). Full details about the wet-lab experimental protocol and computational methods to assemble these genomes are available in the Supplemental Methods. Evaluation of genome properties (N50, contig count, etc.) was performed with QUAST v5.2.0 (Gurevich et al. 2013). Sequencing depth of the read sets was approximated by counting the total number of sequenced bases for each read set using the program composition (https://github.com/richarddurbin/rotate) and dividing that number by the total nucleotide length of the most contiguous genome assembly (*A. calliptera*, 880,445,564 bp). Genome completeness was evaluated with BUSCO v5.5.0 (Benchmarking Universal Single-copy Orthologs) using the “actinopterygii_odb10” dataset from OrthoDB (Manni et al. 2021).

### Graph construction

The cichlid assemblies were integrated into a multiassembly graph using minigraph v0.18-r538 with the graph generation `-xggs` preset. Base alignment `-c` was activated, and the minimum variant length `L` set to the default 50. For the canonical Lake Malawi cichlid graph, the *A. calliptera* fAstCal1.2 assembly was utilized as the backbone, on which the remaining species were integrated to create bubbles of structural variation. Separate graphs were also generated using the seven other Malawi assemblies as backbones for comparison purposes. For each choice of backbone, we generated an additional 30 different random permutations of incorporating species to examine variability in the graph structure and topology. For the canonical version, we also tested values of 1, 2, 5, 10, 25, 100, 250, 500 and 1000 for the minimum variant length `L`. Empirical properties of genome graphs (total nucleotide length, segment and edge count, etc.) were obtained by parsing the GFA output files with custom Python scripts and the `gfatools stat` command in gfatools v0.4-r214 (https://github.com/lh3/gfatools). The `gfatools bubble` algorithm was also used to extract bubble coordinates and properties, which were converted into a BED file for linear visualization in IGV 2.16.1 (Robinson et al. 2011). Genome graphs were also visualized in 2D with Bandage v0.8.1 (Wick et al. 2015).

To determine sequencing coverage over graph segments, we aligned individual non-backbone assemblies onto a graph using minigraph options `-xasm --cov -c` (assembly-to-sequence mapping, coverage, base alignment). The path an assembly traverses through a bubble can only be determined if the sample has “spanning coverage” on the corresponding backbone region in the linear reference bridging from one end of the bubble to the other. The total percentage of backbone nucleotides with spanning coverage by at least one non-backbone assembly was used as an overall, cumulative coverage value, in order to normalise certain statistics (e.g. bubble density) for a fairer comparison between different graphs.

In addition, we generated bi-assembly graphs for fair comparisons of every possible pair of the Lake Malawi cichlid assemblies (8 × 7 = 56). There were two possible graphs for any given pair, because one assembly can act as a backbone to which another (query) was aligned, making this a non-symmetrical comparison. Graph construction `-xgen` and coverage calculation `-xasm --cov` were performed as in subsections above. For each bi-assembly graph, the estimated flexible percentage sequence on the backbone was calculated as how much backbone genomic sequence was located within bubbles, divided by the total backbone length with spanning coverage. Other statistics were calculated by parsing the GFA file or using gfatools v0.4-r214.

### Structural variant calling

The canonical *A. calliptera* backboned graph was utilized for structural variant calling using minigraph with the option `--xasm --call -c`. This allowed the identification of each sample’s allele, or biological path, through every bubble, defined as the sequence formed by the segments in the taken path. We classified structural variant alleles into four main types, following (Crysnanto et al. 2021):

● Complete deletion: only reference sequence present, non-reference has length zero.
● Partial/alternate deletion: reference and non-reference allele present, but non-reference is shorter.
● Partial/alternate insertion: reference and non-reference allele present, but the reference is shorter.
● Complete insertion: only non-reference sequence present, reference has length zero.

Allele lengths were used for intra- and interspecies comparisons using R packages, including heatmaps (ComplexHeatmap v2.14.0, (Gu 2022)), PCA (factoextra v1.0.7, (Kassambara and Mundt 2020)), Pearson correlation (ggcorrplot v0.1.4, (Kassambara 2022)), and the tidyverse (v2.0.0) suite of packages (Wickham et al. 2019). The theoretical complexity of a bubble was determined by `gfatools bubble` as the number of possible paths through the nested segments. Biological paths are denoted as those paths traversed by the assemblies, as determined by minigraph structural variant calling. Bubbles for which the ratio of the shortest divided by the longest biological allele was close to 0 were interpreted as straightforward present-or-absent variation in certain analyses.

### PCR validation and genotyping

Forward and reverse primers were designed for selected bubbles, such that the PCR reaction extended inwards to the bubble, producing different sized products depending on the allele. PCR products were also extracted for Sanger sequencing to confirm their identity. More details are in the Supplemental Methods.

### Genes

The coordinates of structural breakpoints at bubbles were determined according to the *A. calliptera* fAstCal1.2 reference coordinates, which were superimposed against those of gene annotations. For preprocessing, we filtered out certain gene entries as misannotations, removing entries where either: (i) 70% of gene sequence overlap with a single TE fragment, (ii) over 90% overlap with multiple TE fragments, or (iii) annotated with GO term “transposition”. We employed two approaches for SV-to-gene association, the first of which involves computing the overlap or proximity of each individual SV to its closest gene and gene feature. Bubbles that coincide with multiple features were counted only once in decreasing hierarchy: coding exon, 5’-UTR, 3’-UTR, intron, upstream (2 kbp), downstream (2 kbp), and intergenic. For the second, gene-centric approach, we examined each gene by counting the number of SVs in its gene body and flanking 2000 bp regions. Both approaches made use of the bedtools suite (v2.31.0) (Quinlan and Hall 2010) and the GenomicRanges (v1.50.2) + GenomicFeatures (v1.50.4) packages (Lawrence et al. 2013). Gene Ontology functional enrichment analysis for gene sets was performed with the gprofiler2 R package v0.2.2 (Kolberg et al. 2020).

We utilized ODGI v0.7.3 (Guarracino et al. 2022) to estimate the percentage sequence conservation of genes. The minigraph GFA output graph was converted into ODGI’s proprietary format .og with the `build`, `sort` and `chop` functions, after which `odgi pav` was utilized to superimpose gene annotations onto the graph, and compute the percentage sequence conservation of each gene as the number of gene segments traversed by a given sample. We performed this analysis for *A. calliptera* and *M. zebra* backboned graphs with respective Ensembl gene annotations. Gene sequences private to the backbones were investigated further with orthology information from BioMart, Gene Ontology enrichment with gprofiler2 R package v0.2.2 (Kolberg et al., 2020) and protein domain search against the Pfam database Pfam-A.hmm (version 3.1b2, Feb 2015) (Mistry et al. 2021) with hmmsearch.

### Transposable elements

The annotation of transposable elements (TEs) was performed with RepeatModeler v2.0.3 and RepeatMasker v4.1.2-p1, which were included as part of the Dfam TEtools Docker/Singularity container v1.3 (docker://dfam/tetools:1.3) (Storer et al. 2021). We generated de novo repeat families using the `BuildDatabase` and `RepeatModeler` commands, with the `-LTRStruct` option activated. These de novo libraries were then combined with RepBase-derived RepeatMasker libraries (Bao et al. 2015), which were then used to annotate all the genomes with RepeatMasker under options `-e rmblast -no_is`. We overlapped coordinates of TEs with graph bubbles using GenomicRanges (v1.50.2) and GenomicFeatures (v1.50.4). To generate a null model to estimate statistical enrichment of TEs in SVs, we generated 100 random shufflings of SV coordinates using bedtools `shuffle` (v2.31.0) (Quinlan and Hall 2010), with `-incl` parameter to force the coordinates to maintain proximity to the original and be in regions with sequencing coverage.

## Supporting information

CichlidPangenome_Supplemental

## Data Access

Under an Access and Benefit Sharing agreement, these data are made available on an open access basis for research use only. Any person who wishes to use these data for any form of commercial purpose must first enter into a commercial licensing and benefit sharing arrangement with the Government of Malawi.

## Acknowledgements

The Malawi cichlid samples were collected ethically under prescribed permits, and the results and data are published under an Access and Benefit Sharing agreement with the Government of Malawi. We acknowledge the contributions of the employees of Stuart M Grant Ltd, the Malawi Department of Fisheries and the Government of Malawi for their assistance in the collection of samples and the generation of data and results. We also thank Jonathan Price for helpful discussions and insight pertaining to the analysis performed in this paper.

