## Supplementary material for "A pangenomic perspective of the Lake Malawi cichlid radiation reveals extensive structural variation driven by transposable elements": CichlidPangenome_Supplemental

**Supplemental material for:**

### Towards a pangenome view of the Lake Malawi haplochromine cichlid radiation

**<u>Table of Contents</u>**


- **Supplemental Figures**
- **Supplemental Tables**
- **Supplemental Methods**

### Supplemental Figures

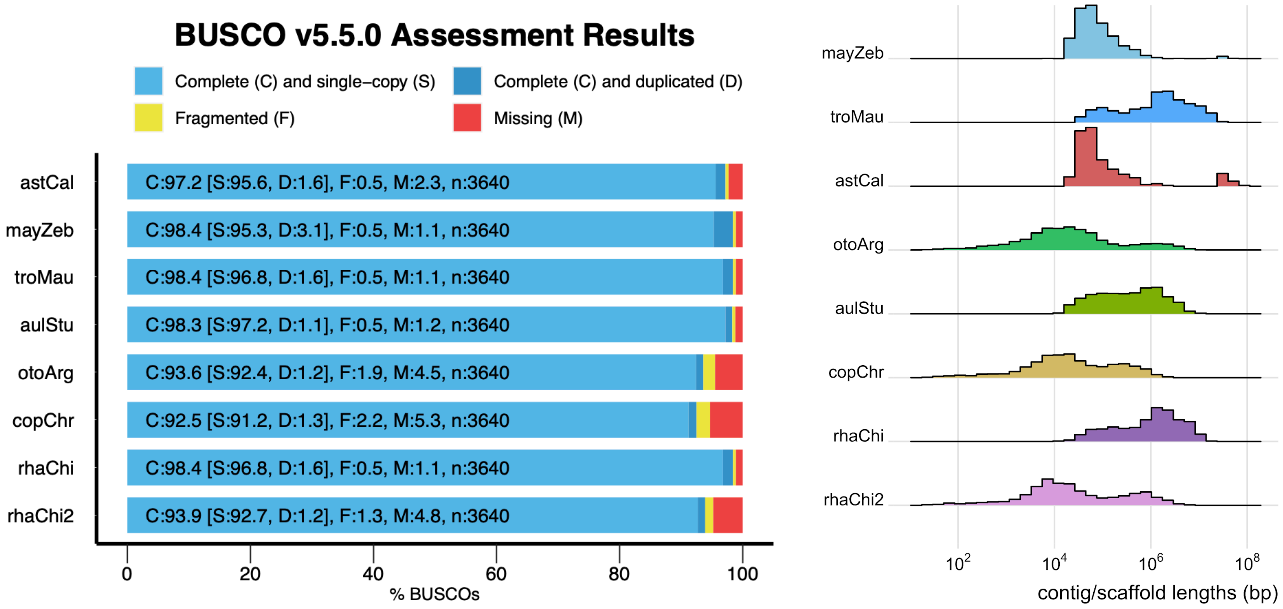


**Supplemental Figure S1.** (Left) Benchmarking Universal Single-copy Orthologs (BUSCO) assessment of genome completeness. Gene completeness was evaluated based on the percentage of detectable genes out of 3,640 essential ray-finned fish genes in the "actinopterygii_odb10" dataset from OrthoDB. (Right) Histogram showing distribution of contig or scaffold sizes for assemblies.

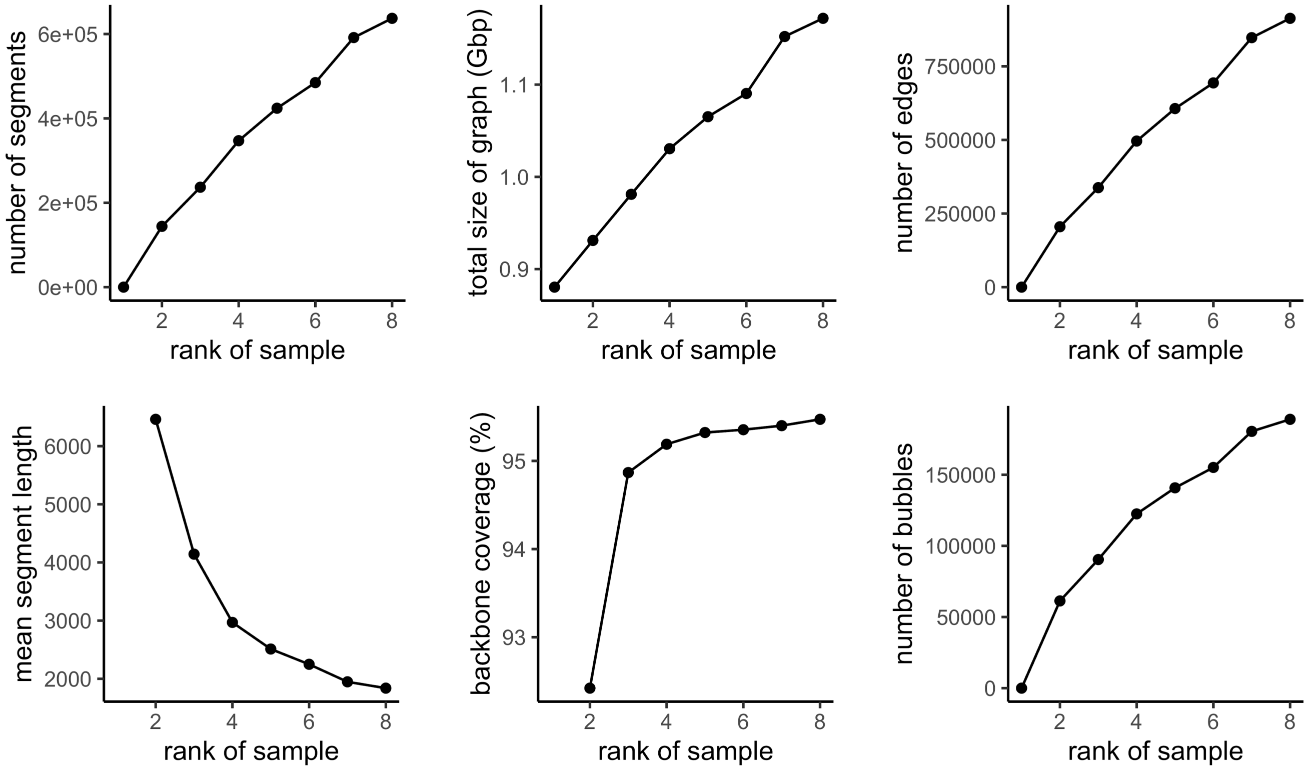


**Supplemental Figure S2.** Growth in the complexity and information content of Lake Malawi cichlid multiassembly graph. Rank of sample (x-axis) denotes the number of assemblies incorporated into the graph, starting at 1 for the *A. calliptera* backbone alone, and rising to 8 as the other seven nonreference assemblies were added.

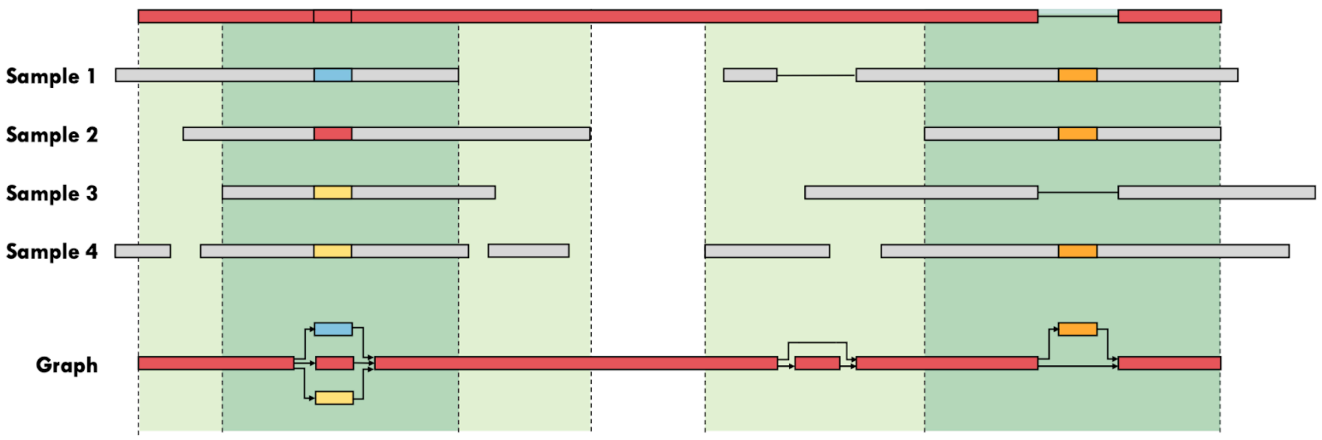


(a) Minigraph algorithm.

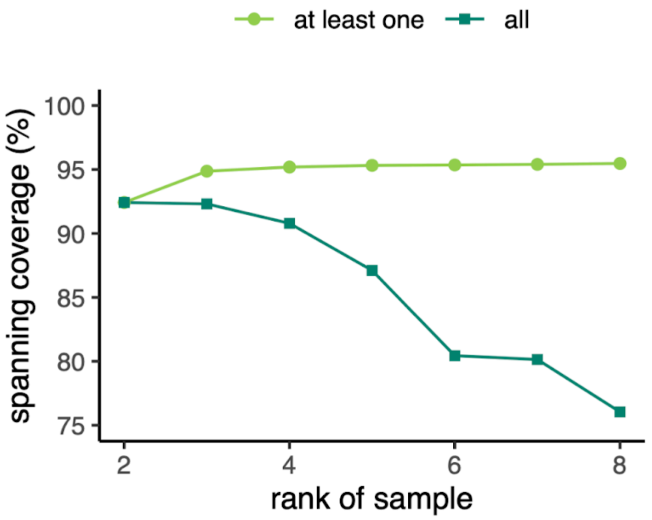


(b) Growth in overall backbone coverage.

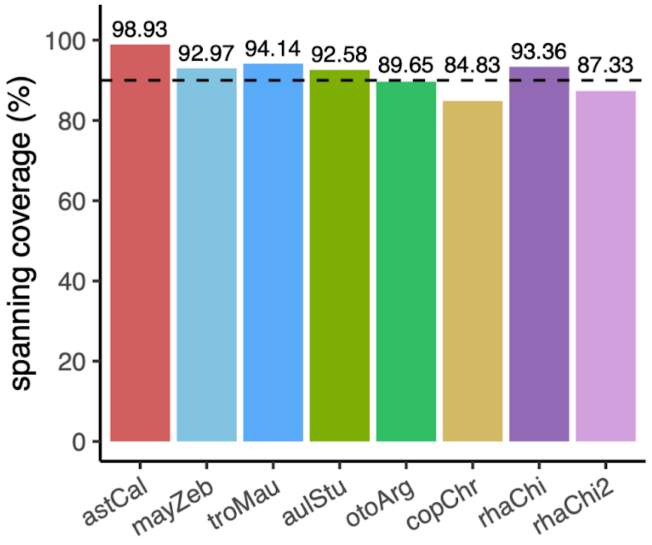


(c) Spanning coverage of individual assemblies.

**Supplemental Figure S3.** Spanning coverage on the *A. calliptera* backbone during the multiassembly graph construction. (a) Illustration of how the minigraph algorithm works, depicting how bubbles are augmented onto the backbone (in red) by aligning the contigs from nonreference assemblies to produce the bottom multiassembly graph. Minigraph does not infer the presence of bubbles at gaps between nonreference contigs. Backbone regions with spanning coverage from at least one sample are colored light green, while those spanned by all are dark green. (b) Growth in the overall spanning coverage on *A. calliptera* fAstCal1.2 backbone as nonreference assemblies were incorporated into the graph. (c) Spanning coverage of individual assemblies when aligned separately on the backbone.

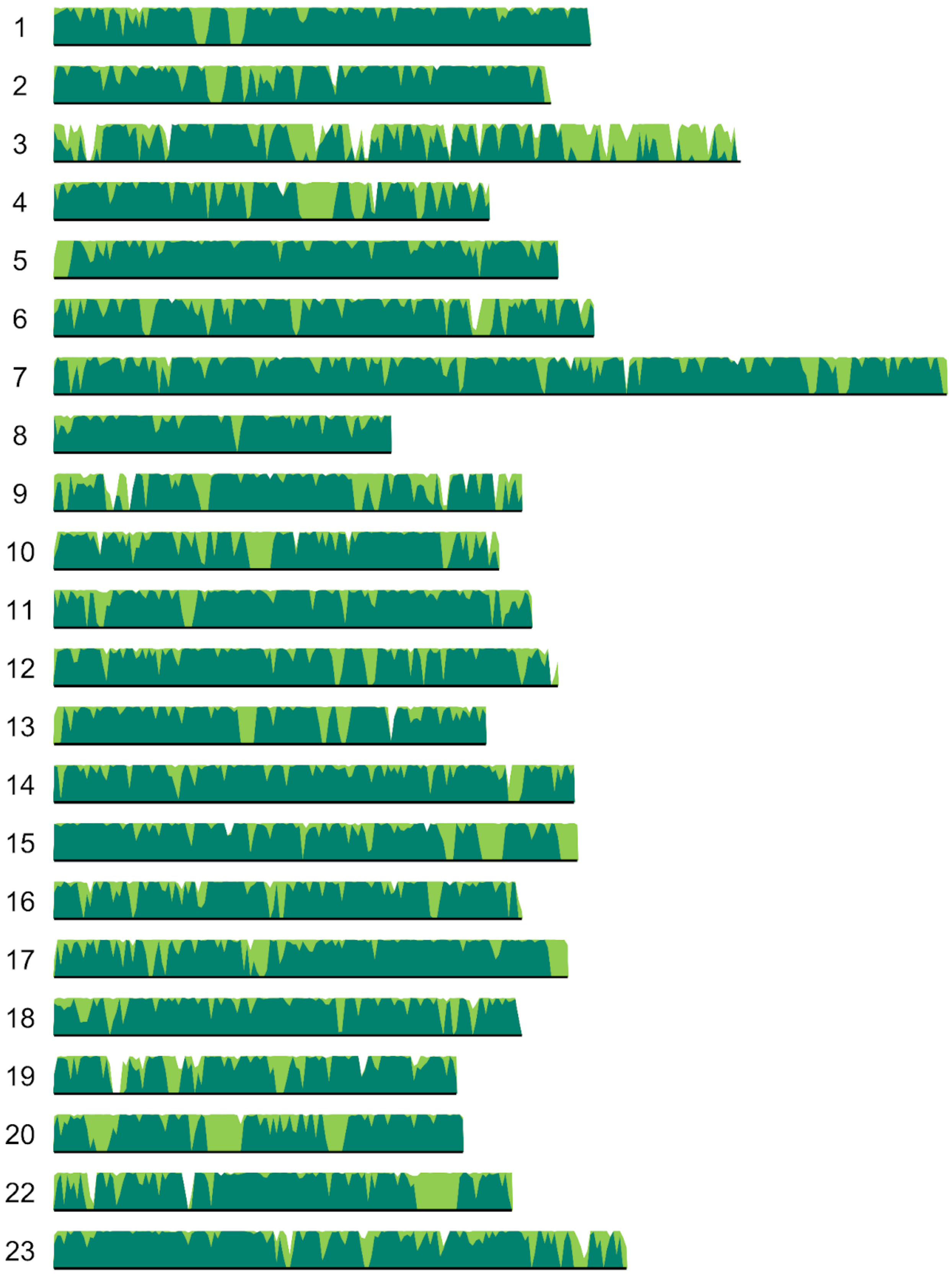


**Supplemental Figure S4.** Spanning coverage across the *Astatotilapia calliptera* fAstCal1.2 backbone assembly, showing 22 linkage groups representing chromosomes. The y-axis denotes sequencing coverage calculated with a window size of 250 bp. Light green denotes regions with coverage from at least one nonreference sample, while dark green denotes coverage from all.

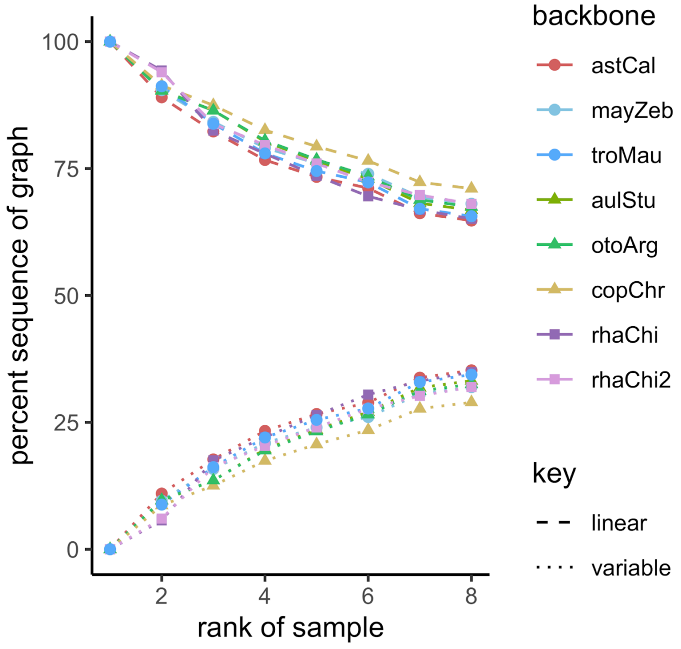


**Supplemental Figure S5.** Relative size of linear and variable components in Lake Malawi cichlid multiassembly graph across different backbone choices.

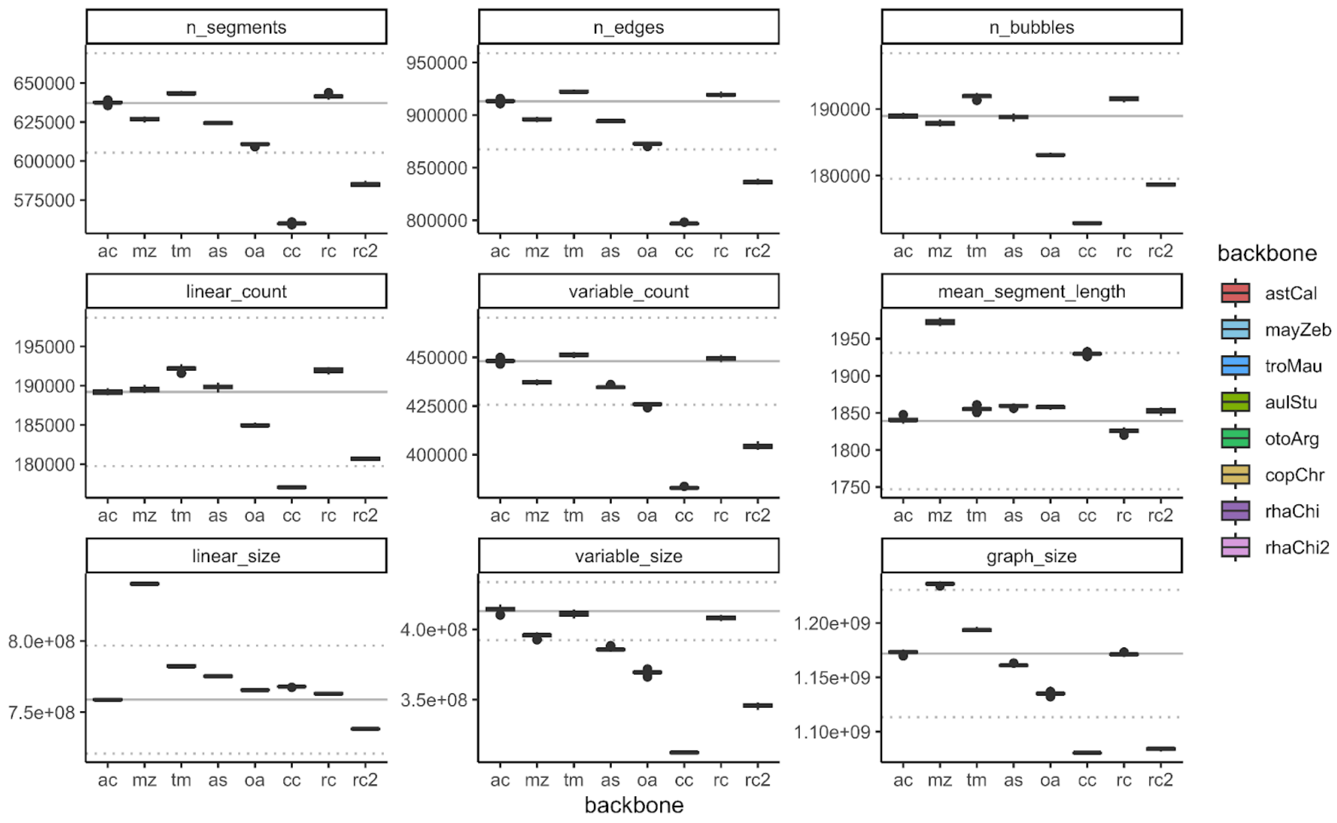


**Supplemental Figure S6:** Stability assessment of Lake Malawi cichlid multiassembly graph across backbone choice and species ordering. Boxplots denote the interquartile range of 30 permutations of incorporating subsequent assemblies for each given backbone in the x-axis. Gray solid lines denote the properties of the canonical graph, with ≈ ranges shown with dotted lines.

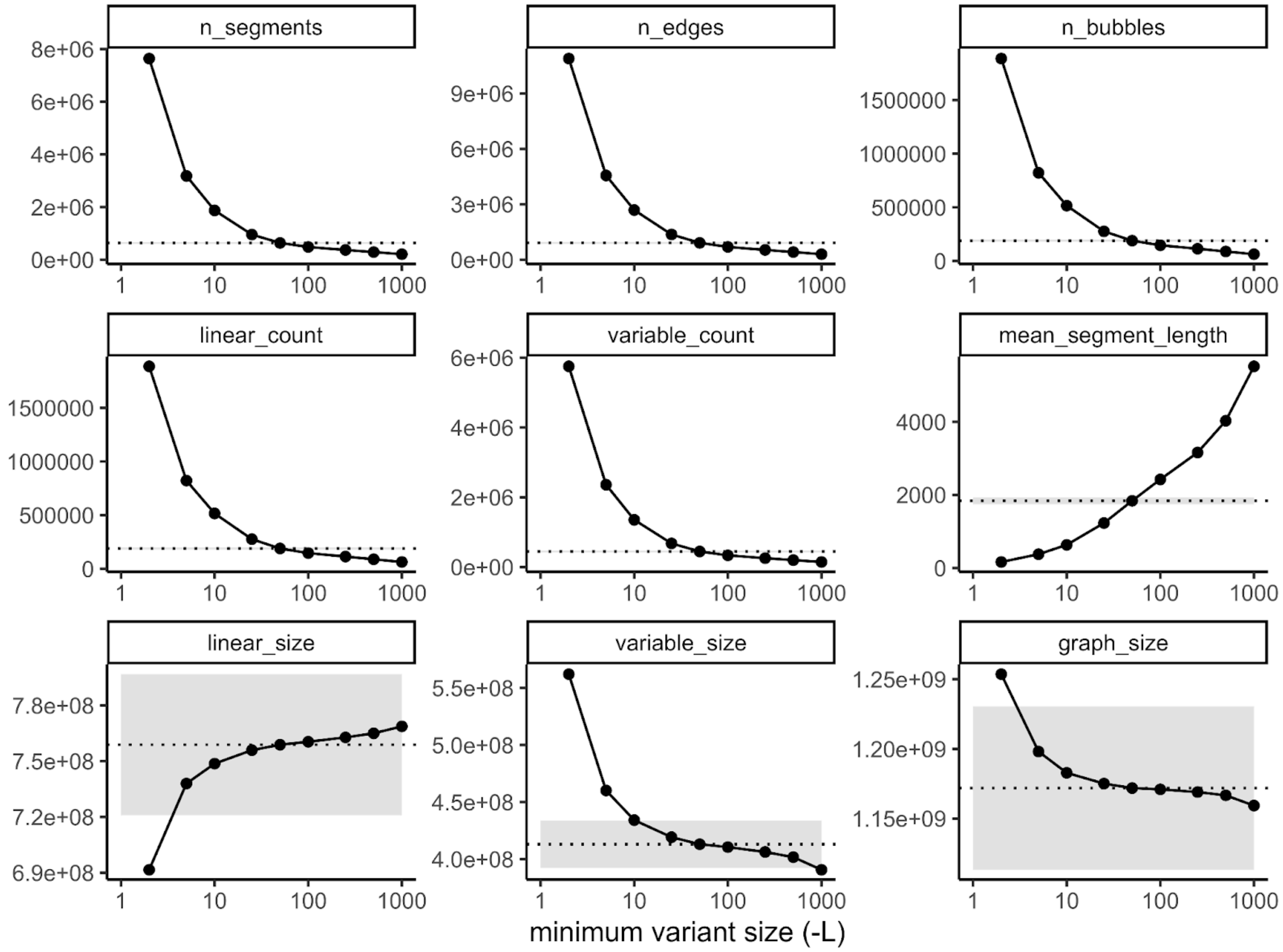


**Supplemental Figure S7.** Stability assessment of Lake Malawi cichlid multiassembly graph across minimum variant length L. Empirical properties were benchmarked across a parameter sweep, of which minigraph's default value L is 50. The canonical *A. calliptera* backbone and ordering of subsequent assemblies was used in all cases. Black dotted lines denote the properties of the default L = 50, with ±5% ranges shaded in gray. The results at L = 1 are omitted because it was an outlier and distorted the visualisation of the current values, but are shown in Supplemental Table S2.

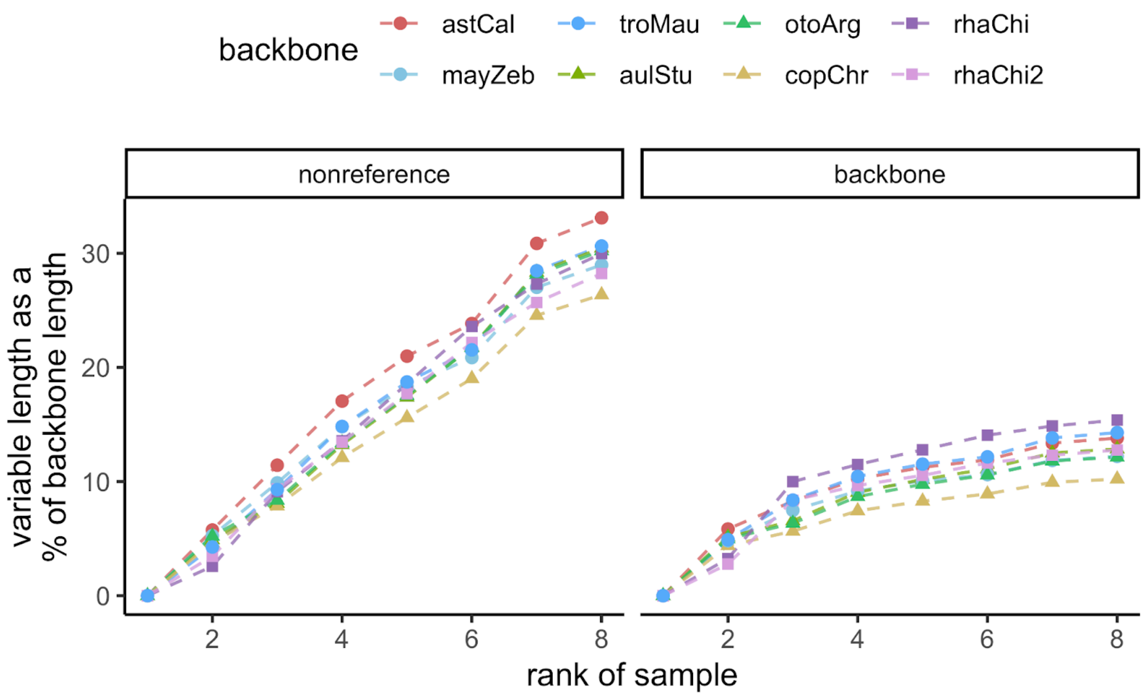


**Supplemental Figure S8.** Growth of the variable component in Lake Malawi multiassembly graph across different backbones. These plots show the amount of sequence that are included in bubbles, based on whether they originated from non-reference assemblies (left) or the backbone (right).

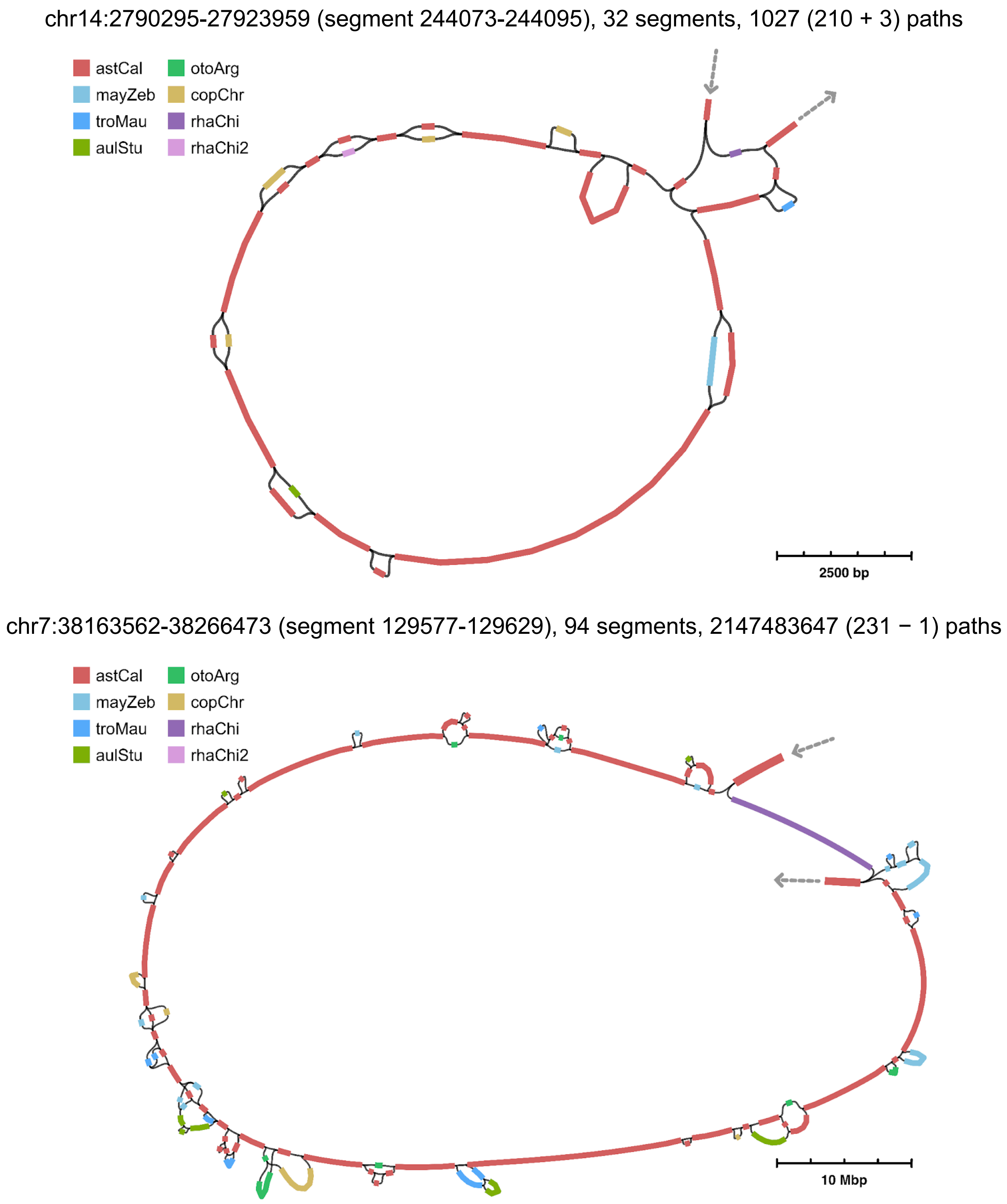


**Supplemental Figure S9:** Intuitive representation of complex structural variation in the multiassembly graph. Graph segments are coloured differently to reflect the original assemblies from which the sequences were derived, with backbone segments coloured red. Arrows denote entry and exit points based on the backbone coordinates.

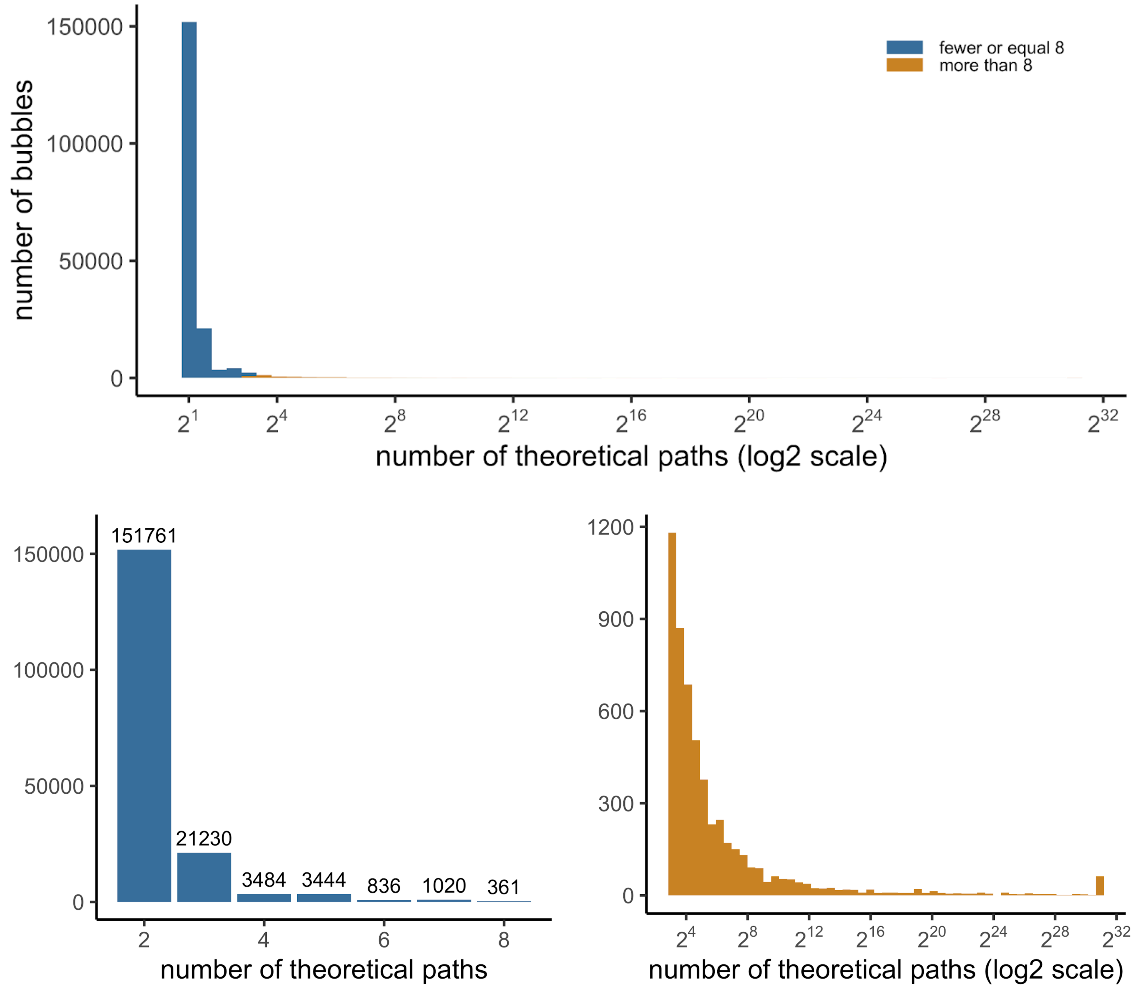


**Supplemental Figure S10.** Theoretical complexity of bubbles in the Lake Malawi cichlid graph. The measure of complexity (x-axis) was obtained by counting the number of theoretical paths for a sample to transverse the bubble.

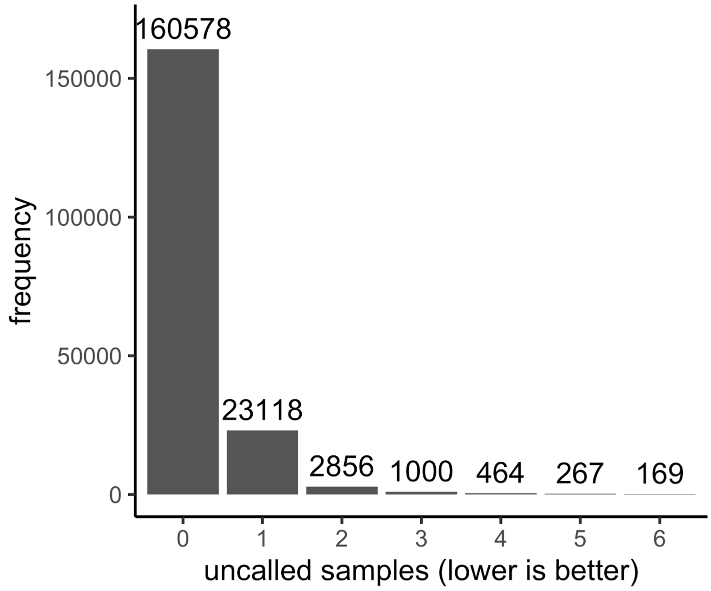


**Supplemental Figure S11.** Success rate of allele calling across 188,452 graph bubbles.

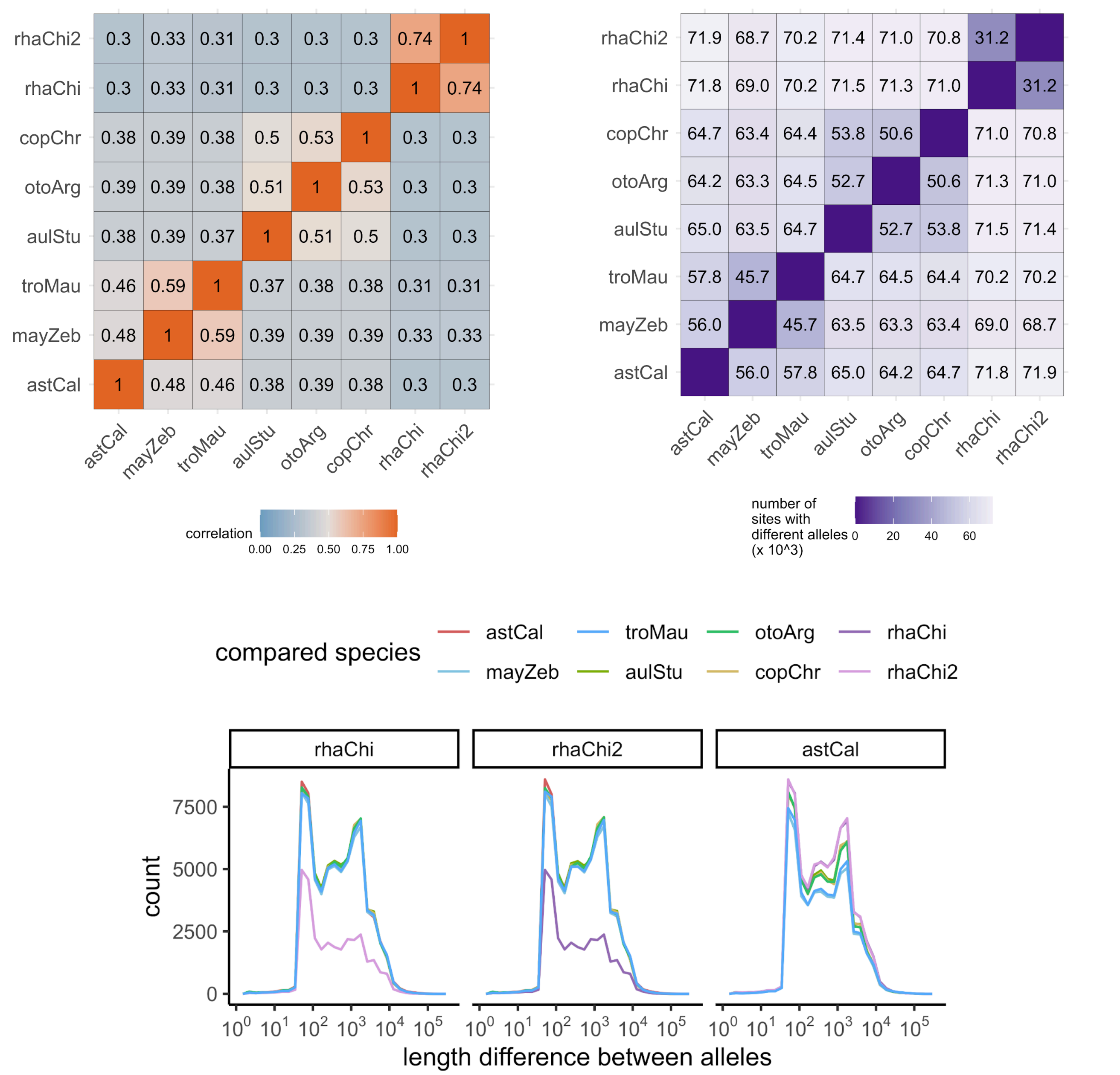


**Supplemental Fig. S12.** Reconstruction of expected relationships between Lake Malawi cichlid assemblies. Analyses were performed for the 160,572 bubbles where there was complete allelic information for all the samples. The two *Rhamphochromis* individuals were closer in distance across all measures.

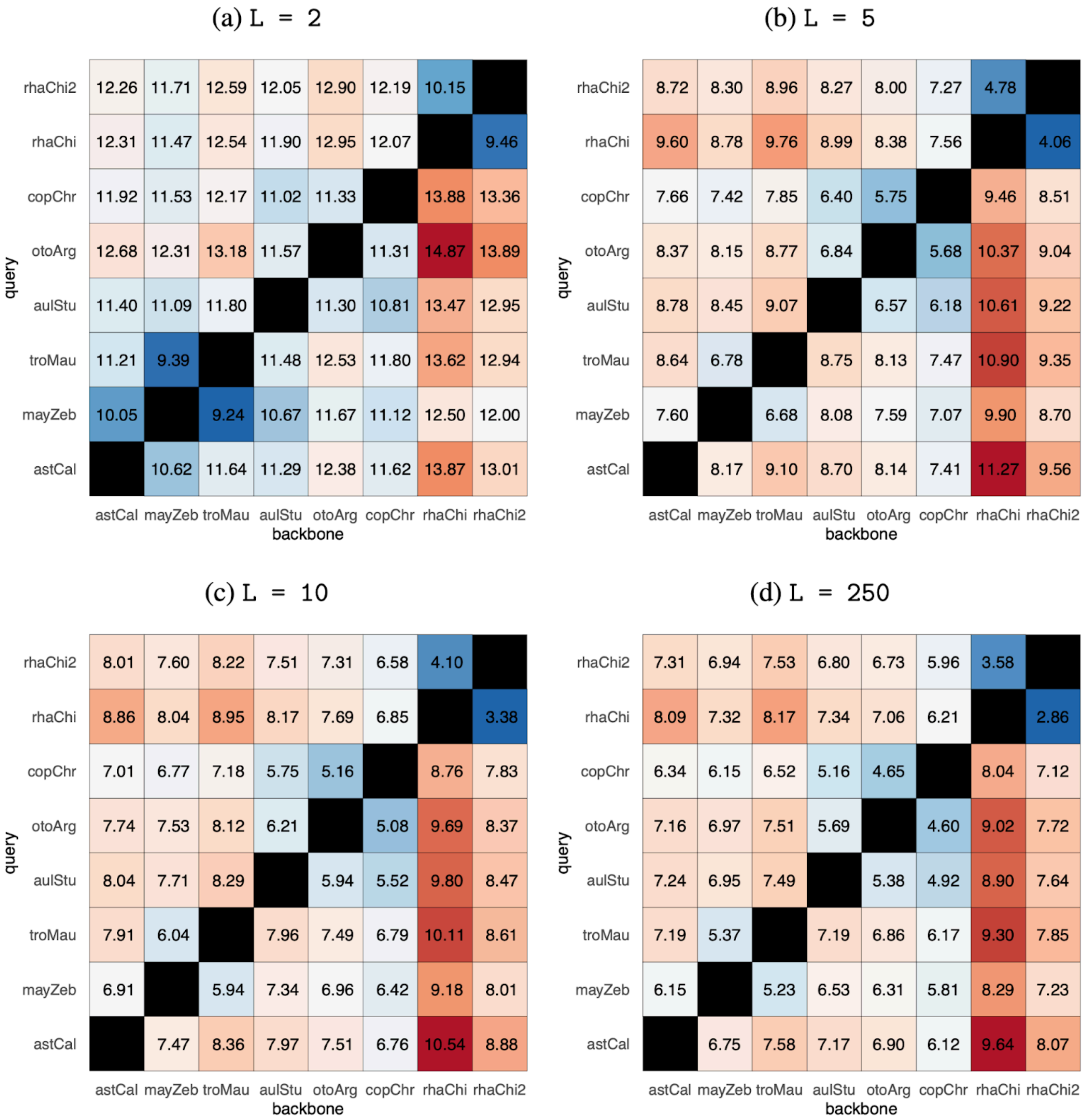


**Supplemental Figure S13.** Estimated percentage sequence of structural variants across parameter sweep of minimum variant size L. Values were calculated using bi-assembly graphs built from pairs of the Lake Malawi haplochromine cichlid assemblies. For each possible pair, the sequences of a query (y-axis) were aligned onto a backbone (x-axis), and the percentage of flexible sequence on the backbone was estimated based on how much of these backbone regions with spanning coverage was located inside bubbles.

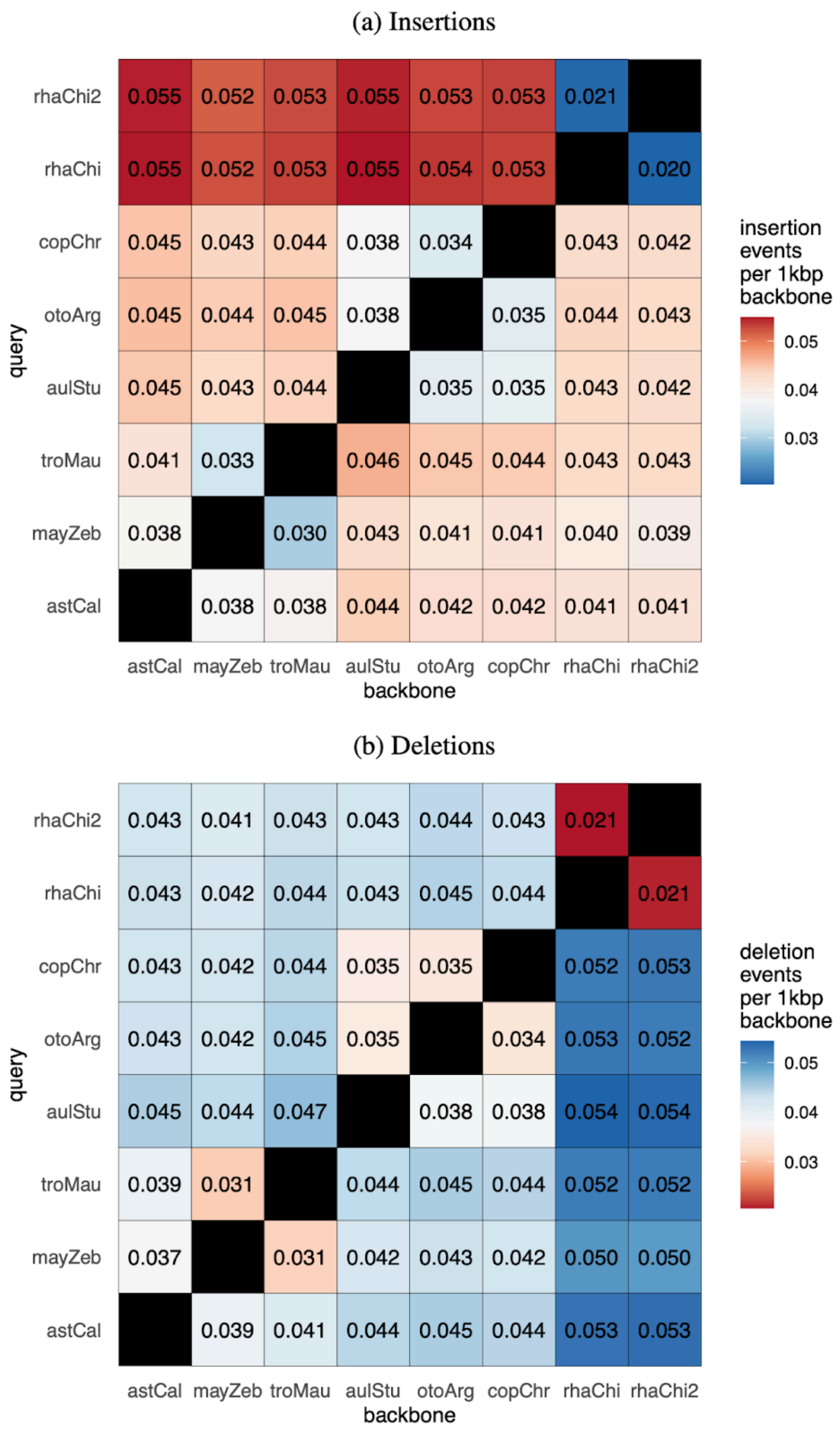


**Supplemental Fig. S14.** Insertion and deletion densities estimated from bi-assembly graphs between pairs of cichlid assemblies. For each possible pair, the sequences of a query (y-axis) were aligned onto a backbone (x-axis), and the density was estimated based on how many events are discovered per 1 kbp of backbone regions with spanning coverage. Minimum variant size L set to 50.

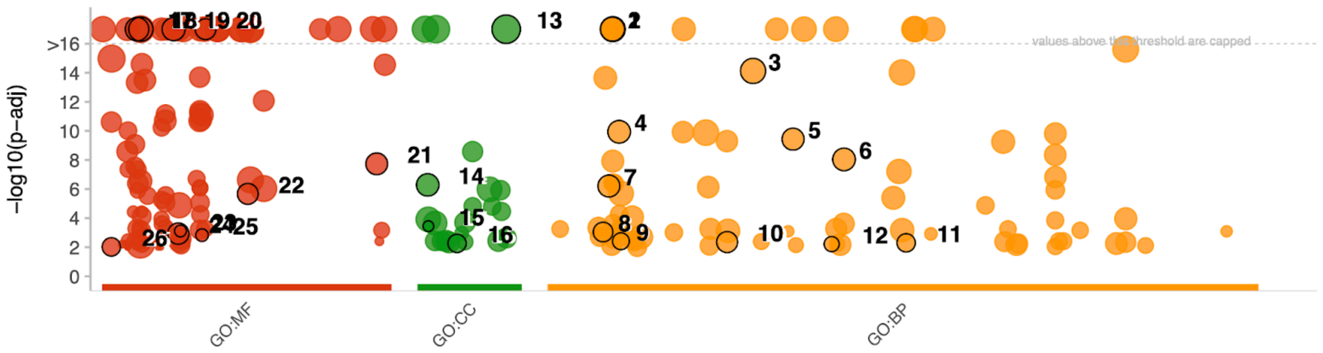


| id | source | term_id | term_name | term_size | intersection_size | | p_value |
|---|---|---|---|---|---|---|---|
| 1 | GO:BP | GO:0006810 | transport | 2275 | 1727 | | 8.0e−35 |
| 2 | GO:BP | GO:0006793 | phosphorus metabolic process | 1435 | 1110 | | 3.7e−26 |
| 3 | GO:BP | GO:0032502 | developmental process | 2524 | 1809 | | 7.5e−15 |
| 4 | GO:BP | GO:0007166 | cell surface receptor signaling pathway | 768 | 586 | | 1.1e−10 |
| 5 | GO:BP | GO:0035556 | intracellular signal transduction | 595 | 462 | | 3.7e−10 |
| 6 | GO:BP | GO:0044281 | small molecule metabolic process | 763 | 575 | | 9.1e−09 |
| 7 | GO:BP | GO:0006629 | lipid metabolic process | 572 | 434 | | 6.2e−07 |
| 8 | GO:BP | GO:0006325 | chromatin organization | 236 | 185 | | 9.0e−04 |
| 9 | GO:BP | GO:0007268 | chemical synaptic transmission | 97 | 82 | | 3.9e−03 |
| 10 | GO:BP | GO:0030030 | cell projection organization | 389 | 290 | | 4.4e−03 |
| 11 | GO:BP | GO:0050790 | regulation of catalytic activity | 153 | 123 | | 5.0e−03 |
| 12 | GO:BP | GO:0043087 | regulation of GTPase activity | 62 | 55 | | 6.1e−03 |
| 13 | GO:CC | GO:0110165 | cellular anatomical entity | 14671 | 9469 | | 2.4e−32 |
| 14 | GO:CC | GO:0005856 | cytoskeleton | 716 | 528 | | 5.2e−07 |
| 15 | GO:CC | GO:0005891 | voltage−gated calcium channel complex | 28 | 28 | | 3.6e−04 |
| 16 | GO:CC | GO:0034702 | monoatomic ion channel complex | 147 | 116 | | 5.5e−03 |
| 17 | GO:MF | GO:0005215 | transporter activity | 1163 | 933 | | 5.4e−38 |
| 18 | GO:MF | GO:0005524 | ATP binding | 1621 | 1252 | | 1.6e−36 |
| 19 | GO:MF | GO:0016301 | kinase activity | 920 | 729 | | 4.4e−26 |
| 20 | GO:MF | GO:0030695 | GTPase regulator activity | 356 | 301 | | 1.1e−17 |
| 21 | GO:MF | GO:0140657 | ATP−dependent activity | 537 | 406 | | 1.8e−08 |
| 22 | GO:MF | GO:0042578 | phosphoric ester hydrolase activity | 369 | 282 | | 2.1e−06 |
| 23 | GO:MF | GO:0016849 | phosphorus−oxygen lyase activity | 33 | 32 | | 7.7e−04 |
| 24 | GO:MF | GO:0016746 | acyltransferase activity | 436 | 317 | | 1.2e−03 |
| 25 | GO:MF | GO:0030215 | semaphorin receptor binding | 37 | 35 | | 1.5e−03 |
| 26 | GO:MF | GO:0003774 | cytoskeletal motor activity | 156 | 121 | | 9.5e−03 |

g:Profiler (biit.cs.ut.ee/gprofiler)

**Supplemental Figure S15.** Functional enrichment analysis of genes containing structural variants, with non-redundant driver terms as highlighted by g:Profiler.

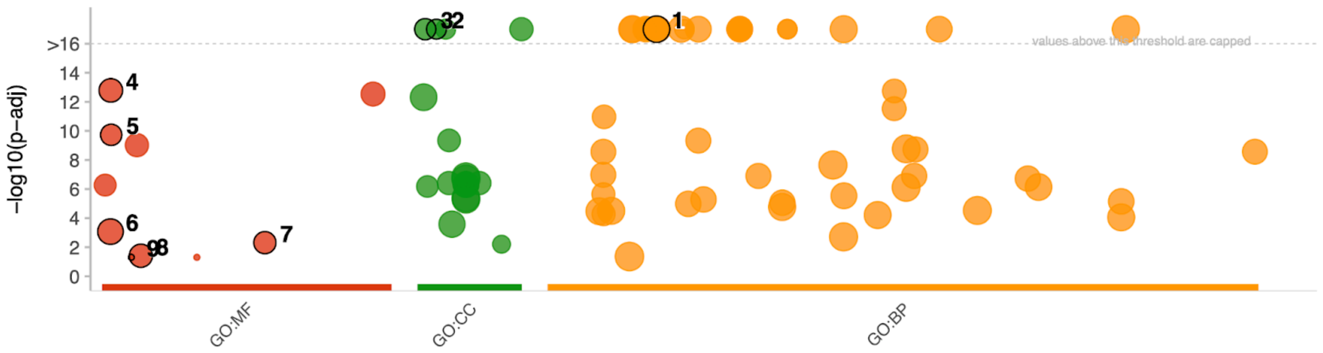


| id | source | term_id | term_name | term_size | intersection_size | | p_value |
|---|---|---|---|---|---|---|---|
| 1 | GO:BP | GO:0010467 | gene expression | 2640 | 628 | | 4.4e−40 |
| 2 | GO:CC | GO:0016442 | RISC complex | 174 | 104 | | 9.2e−38 |
| 3 | GO:CC | GO:0005730 | nucleolus | 277 | 110 | | 8.3e−20 |
| 4 | GO:MF | GO:0003700 | DNA−binding transcription factor activity | 706 | 179 | | 1.6e−13 |
| 5 | GO:MF | GO:0003735 | structural constituent of ribosome | 290 | 88 | | 1.9e−10 |
| 6 | GO:MF | GO:0003677 | DNA binding | 1711 | 306 | | 8.5e−04 |
| 7 | GO:MF | GO:0046983 | protein dimerization activity | 399 | 87 | | 4.9e−03 |
| 8 | GO:MF | GO:0005525 | GTP binding | 600 | 117 | | 3.9e−02 |
| 9 | GO:MF | GO:0004860 | protein kinase inhibitor activity | 17 | 9 | | 5.0e−02 |

g:Profiler (biit.cs.ut.ee/gprofiler)

**Supplemental Figure S16.** Functional enrichment analysis of genes without structural variants in the gene body and 2000 bp upstream or downstream, with non-redundant driver terms as highlighted by g:Profiler.

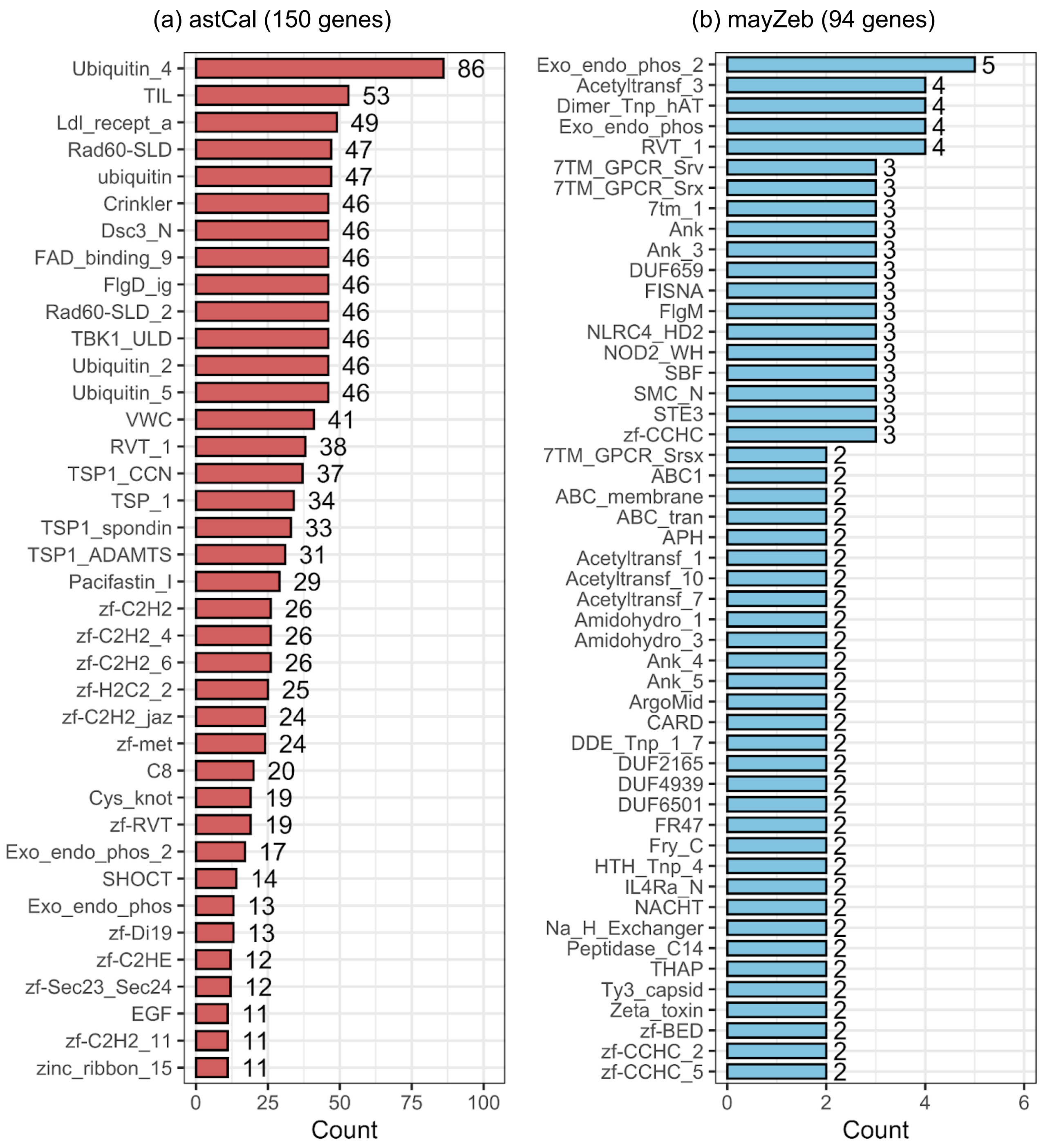


**Supplemental Figure S17.** Pfam domain annotation for private genes in *A. calliptera* and *M. zebra*. Domain detection achieved by alignment of gene sequences against the Pfam database with HMMER 3.3.

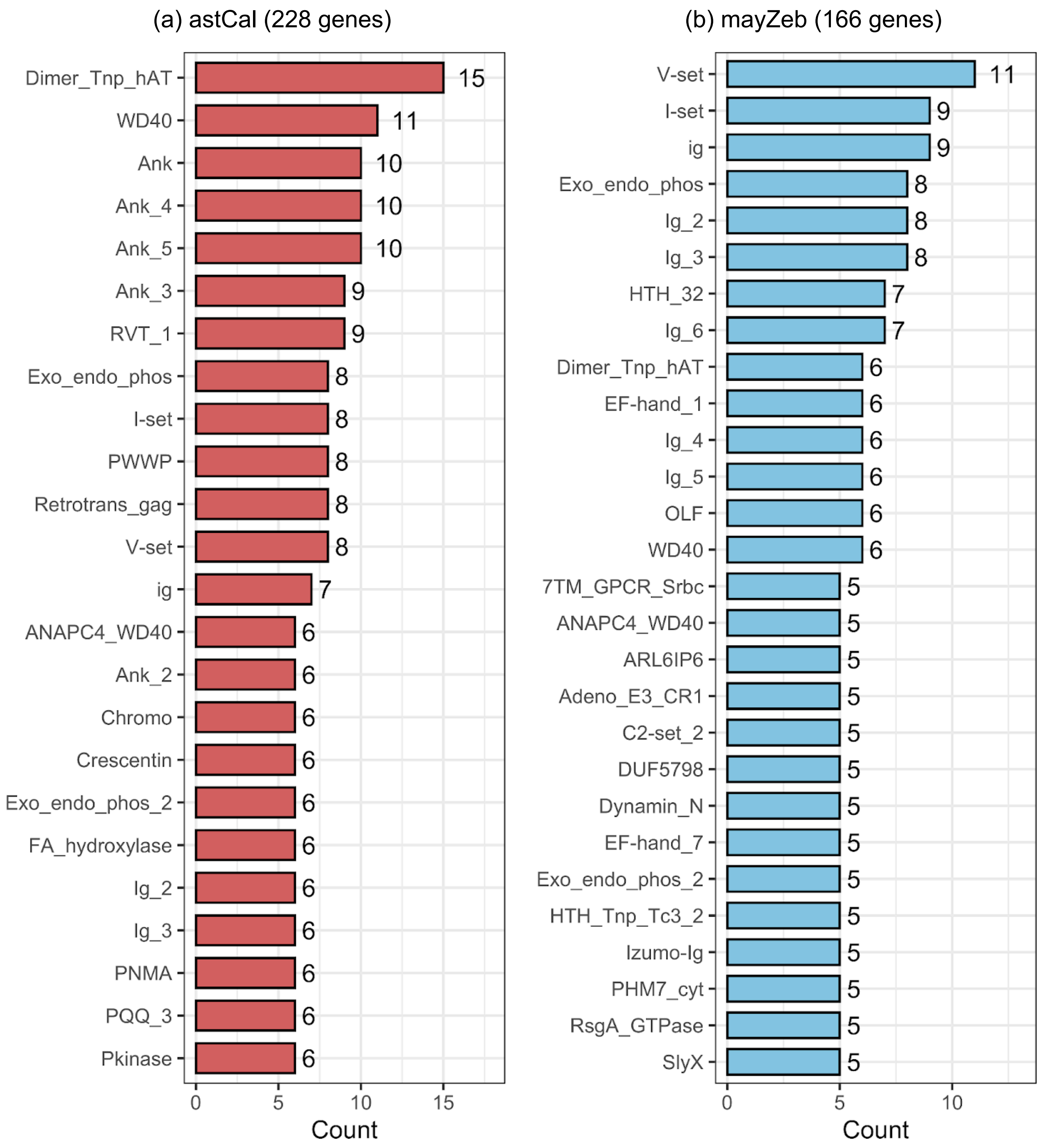


**Supplemental Figure S18.** Pfam domain annotation for genes mistakened to be private to *A. calliptera* and *M. zebra*. Domain detection achieved by alignment of gene sequences against the Pfam database with HMMER 3.3.

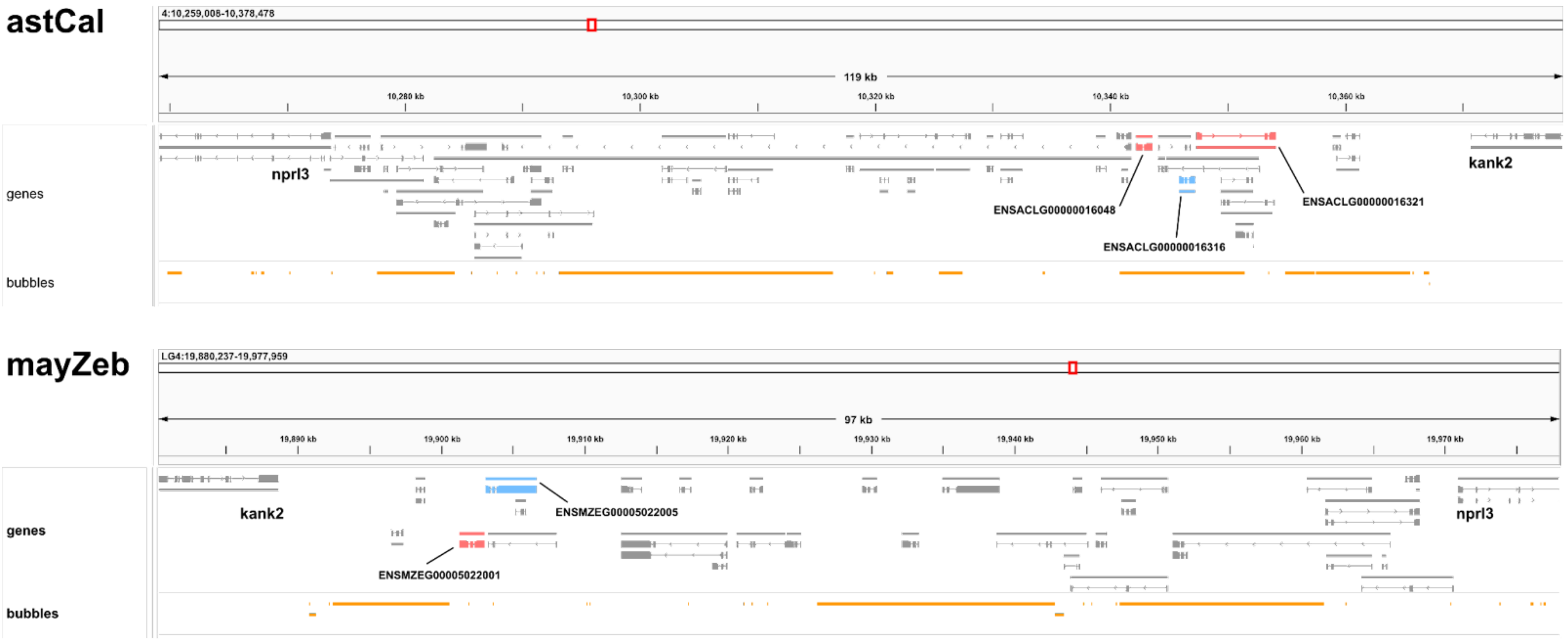


**Supplemental Figure S19.** Inaccurate structural variant detection at the haemoglobin MN locus caused by backbone assembly errors. Globin genes appear to be poorly assembled in the *A. calliptera* fAstCal1.2 genome (top), compared to in *Maylandia zebra*. The labeled globin alpha-B (red) and beta-A (blue) subunits in *A. calliptera* were determined by ODGI to be unique to the backbone, but actually had matching orthologs in *M. zebra* based on BioMart. Note that the orientation of the chromosomes are reversed.

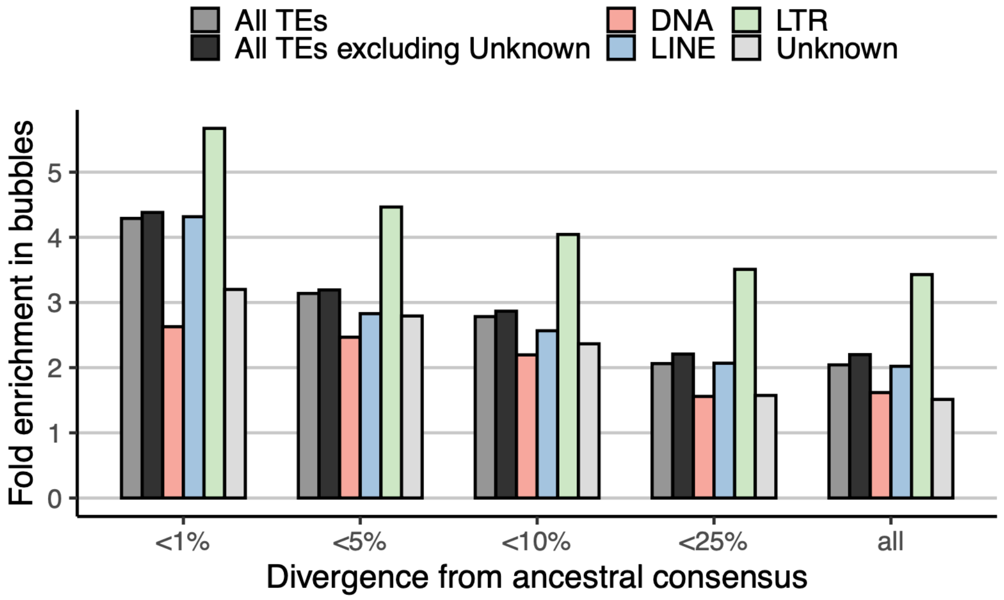


**Supplemental Figure S20.** Transposon enrichment across different sequence divergence thresholds. SINEs, Helitrons and Retroposons are not shown because their overly small values make the calculation unreliable.

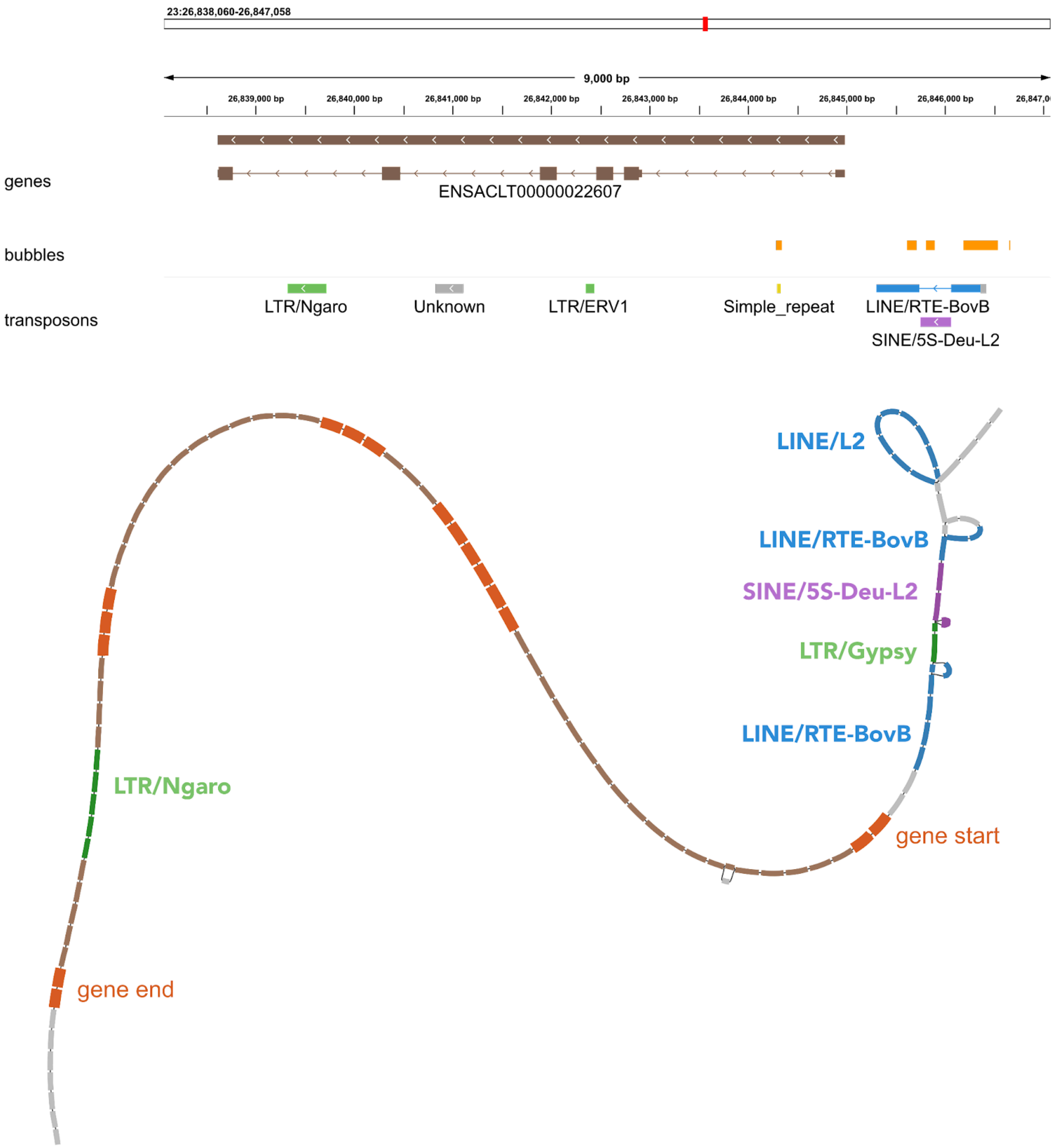


**Supplemental Figure S21.** TE insertions around the *fhl2b* gene, showing the conserved SINE element in purple. Segments are coloured manually based on the presence of TEs. Exons and introns are coloured dark orange and brown respectively, with the former having a thicker width.

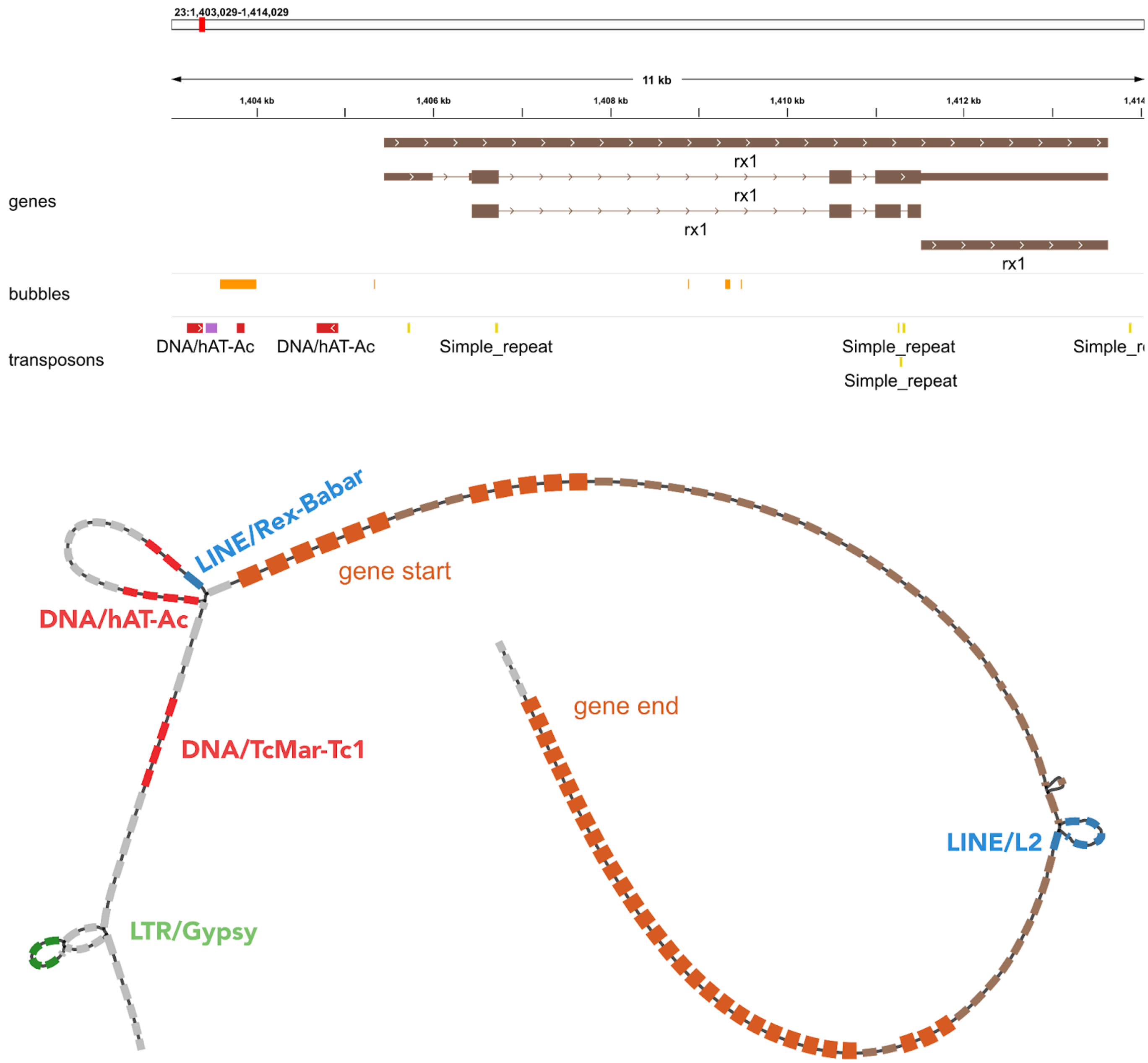


**Supplemental Figure S22.** E insertions around the *rx1* gene, showing the conserved SINE element in purple. Segments are coloured manually based on the presence of TEs. Exons and introns are coloured dark orange and brown respectively, with the former having a thicker width. The first bubble from the left is known from previous research, hosting three possible alleles: complete deletion, partial insertion of 413 bp and the full length of 831 bp, each of which contribute to different opsin palettes in cichlid vision.

### Supplemental Tables

| | astCal | mayZeb | troMau | aulStu | otoArg | copChr | rhaChi | rhaChi2 |
|---|---|---|---|---|---|---|---|---|
| **segment count** | 637,237 | 626,826 | 642,989 | 625,018 | 610,555 | 559,685 | 642,806 | 587,152 |
| **edge count** | 913,087 | 896,071 | 921,610 | 895,007 | 872,224 | 796,680 | 921,326 | 839,389 |
| **mean segment length** | 1,839.04 | 1,970.12 | 1,854.30 | 1,857.58 | 1,858.03 | 1,930.61 | 1,823.03 | 1,847.71 |
| **graph size, Gbp** | 1.172 | 1.235 | 1.192 | 1.161 | 1.134 | 1.081 | 1.172 | 1.085 |
| **linear percentage** | 64.75% | 68.07% | 65.61% | 66.80% | 67.47% | 71.05% | 65.10% | 68.03% |
| **variable percentage** | 35.25% | 31.93% | 34.39% | 33.20% | 32.53% | 28.95% | 34.90% | 31.94% |
| **backbone coverage** | 95.47% | 89.26% | 94.70% | 95.09% | 93.06% | 89.68% | 97.15% | 94.94% |
| **additional bases** | 33.11% | 28.98% | 30.64% | 30.46% | 30.22% | 26.36% | 29.98% | 28.23% |
| **flexible on backbone** | 13.81% | 12.21% | 14.29% | 12.86% | 12.14% | 10.22% | 15.39% | 12.76% |
| **bubble count** | 188,944 | 188,006 | 192,055 | 189,201 | 183,423 | 172,846 | 191,963 | 178,693 |
| **bubble density, per kbp** | 0.2146 | 0.1964 | 0.2104 | 0.2126 | 0.2106 | 0.2021 | 0.2129 | 0.2112 |
| **bubble density, per kbp (coverage corrected)** | 0.2248 | 0.2200 | 0.2222 | 0.2236 | 0.2263 | 0.2254 | 0.2192 | 0.2225 |

**Supplemental Table S1.** Empirical properties of Lake Malawi cichlid multiassembly graphs constructed with different backbones. Linear and variable percentages are with respect to the entire graph. Backbone coverage denotes percentage sequence with at least spanning coverage with one aligned assembly. Additional bases is the percentage of extra bases relative to the size of the backbone.

| | 1 | 2 | 5 | 10 | 25 | 50 | 100 | 250 | 500 | 1000 |
|---|---|---|---|---|---|---|---|---|---|---|
| **segment count** | 2,446 | 7,638,583 | 3,183,041 | 1,870,374 | 955,101 | 637,237 | 483,066 | 370,077 | 289,658 | 210,137 |
| **edge count** | 3,085 | 10,893,890 | 4,564,107 | 2,689,473 | 1,368,504 | 913,087 | 693,044 | 531,467 | 415,489 | 300,430 |
| **mean segment length** | 403618.03 | 164.11 | 376.40 | 632.40 | 1230.37 | 1839.04 | 2424.06 | 3158.94 | 4027.89 | 5517.15 |
| **graph size, Gbp** | 0.987 | 1.254 | 1.198 | 1.183 | 1.175 | 1.172 | 1.171 | 1.169 | 1.167 | 1.159 |
| **linear percentage** | 81.77% | 55.17% | 61.60% | 63.30% | 64.33% | 64.75% | 64.94% | 65.25% | 65.57% | 66.31% |
| **variable percentage** | 18.23% | 44.83% | 38.40% | 36.70% | 35.67% | 35.25% | 35.06% | 34.75% | 34.43% | 33.69% |
| **additional bases** | 12.13% | 42.38% | 36.08% | 34.35% | 33.47% | 33.11% | 33.00% | 32.78% | 32.52% | 31.68% |
| **flexible on backbone** | 8.31% | 21.45% | 16.18% | 14.96% | 14.14% | 13.81% | 13.63% | 13.36% | 13.11% | 12.69% |
| **bubble count** | 417 | 1,887,173 | 822,148 | 515,500 | 275,825 | 188,944 | 145,915 | 113,700 | 88,162 | 63,004 |
| **bubble density, per kbp** | 0.0005 | 2.1435 | 0.9338 | 0.5855 | 0.3133 | 0.2146 | 0.1657 | 0.1291 | 0.1001 | 0.0716 |

**Supplemental Table S2.** Stability assessment of Lake Malawi cichlid multiassembly graph across minimum variant length L. Empirical properties were benchmarked across a parameter sweep, of which minigraph's default value for L is 50. The canonical *A. calliptera* backbone and ordering of subsequent assemblies was used in all cases. Setting L = 1 resulted in an overly simple graph, which was likely an outlier behaviour in minigraph.

(a) Bubbles with singleton SVs

| Bubble | Nearest Gene | Ensembl ID | Gene Description | Score | Heterozygotes | Sanger Confirmation |
|---|---|---|---|---|---|---|
| 3:9790027-9790027 | rpgrip1 | ENSACLG00000020186 | RPGR interacting protein 1 | 5/5 | - | Complete |
| 20:15022145-15023430 | klhl7 | ENSACLG00000006581 | kelch-like protein 7 | 5/5 | - | Complete |
| 23:12757935-12758244 | tprg1 | ENSACLG00000024576 | tumor protein p63-regulated gene 1-like protein | 5/5 | - | Complete |
| 7:42326247-42326247 | anpepb | ENSACLG00000017196 | aminopeptidase Ey-like | 5/5 | - | Complete |
| 5:33820642-33820642 | mfsd4aa | ENSACLG00000025203 | major facilitator superfamily domain containing 4A | 5/5 | 1/5 (T) | Complete |
| 15:16298262-16298895 | dtnba | ENSACLG00000027349 | dystrobrevin, beta a | 5/5 | 1/5 (O) | 2/5 confirmed |
| 5:30881965-30882447 | mitfa | ENSACLG00000022427 | melanocyte inducing transcription factor a | 5/5 | 2/5 (O,C) | 4/5 confirmed |
| 16:5354683-5354683 | lcp1 | ENSACLG00000015018 | lymphocyte cytosolic protein 1 | 4/5 (C) | - | Complete |

(b) Bubbles with more common SVs

| Bubble | Nearest Gene | Ensembl ID | Gene Description | Score | Heterozygotes | Sanger Confirmation |
|---|---|---|---|---|---|---|
| 10:13464861-13465174 | kcnab3 | ENSACLG00000017346 | voltage-gated potassium channel subunit beta-3-like | 5/5 | - | Complete |
| 7:51031051-51031051 | - | ENSACLG00000002767 | coiled-coil-helix-coiled-coil-helix domain containing 3b | 5/5 | 2/5 (O,R♀) | Complete |
| 15:25833370-25833370 | adam17b | ENSACLG00000014403 | ADAM metallopeptidase domain 17 | 5/5 | - | 4/5 confirmed |
| 10:2382458-2382689 | dgat2 | ENSACLG00000026652 | diacylglycerol O-acyltransferase 2 | 5/5 | - | 3/5 confirmed |
| 7:16423102-16423419 | ptges | ENSACLG00000027791 | prostaglandin E synthase | 5/5 | - | 2/5 confirmed |
| 3:41434172-41435346 | - | ENSACLG00000021983 | H-2 class II histocompatibility antigen, E-S beta chain-like | 4/5 (O) | - | Complete |
| 10:24736910-24737125 | nsd1b | ENSACLG00000015768 | histone-lysine N-methyltransferase, H3 lysine-36 and H4 lysine-20 specific-like | 4/5 (C) | - | Complete |
| 9:28296874-28297388 | gabrr3b | ENSACLG00000013807 | gamma-aminobutyric acid receptor subunit rho-3-like | 4/5 (C) | - | 3/5 confirmed |

**Supplemental Table S3.** Summary of PCR validation results on original samples. The score column indicates the number of samples with PCR bands that matched their predicted allele lengths from the graph. Individuals with a band not of the expected size are indicated in parentheses, as well as those that exhibited heterozygosity. Sanger sequencing has been attempted to check the amplified PCR products. O = *Otopharynx argyrosoma*, C = *Copadichromis chrysonotus*, R = *Rhamphochromis sp. "chilingali"*.

(a) *A. calliptera* backbone, universal reliable genes: 20,785

| | Reliable genes for PAV analysis | >90% conserved | | >90% conserved, reliable for all samples | |
|---|---|---|---|---|---|
| | | gene body | exon | gene body | exon |
| **astCal** | 25,880 | 25,864 (99.9%) | 25,876 (100.0%) | 20,959 (99.9%) | 20,969 (100.0%) |
| **mayZeb** | 24,676 | 22,152 (89.8%) | 23,433 (95.0%) | 18,992 (90.6%) | 20,029 (95.5%) |
| **troMau** | 24,784 | 22,285 (89.9%) | 23,549 (95.0%) | 19,036 (90.8%) | 20,027 (95.5%) |
| **aulStu** | 24,551 | 21,817 (88.9%) | 23,255 (94.7%) | 18,812 (89.7%) | 19,961 (95.2%) |
| **otoArg** | 23,903 | 21,296 (89.1%) | 22,627 (94.7%) | 18,840 (89.8%) | 19,933 (95.0%) |
| **copChr** | 23,123 | 20,633 (89.2%) | 21,882 (94.6%) | 18,802 (89.6%) | 19,910 (94.9%) |
| **rhaChi** | 24,636 | 21,715 (88.1%) | 23,210 (94.2%) | 18,722 (89.3%) | 19,894 (94.9%) |
| **rhaChi2** | 23,354 | 20,627 (88.3%) | 21,997 (94.2%) | 18,681 (89.1%) | 19,849 (94.6%) |

(b) *M. zebra* backbone, universal reliable genes: 20,266

| | Reliable genes for PAV analysis | >90% conserved | | >90% conserved, reliable for all samples | |
|---|---|---|---|---|---|
| | | gene body | exon | gene body | exon |
| **astCal** | 24,047 | 21,887 (91.0%) | 23,180 (96.4%) | 18,612 (91.8%) | 19,641 (96.9%) |
| **mayZeb** | 27,343 | 27,308 (99.9%) | 27,330 (100.0%) | 20,236 (99.9%) | 20,254 (99.9%) |
| **troMau** | 24,692 | 22,990 (93.1%) | 23,932 (96.9%) | 19,027 (93.9%) | 19,746 (97.4%) |
| **aulStu** | 24,316 | 21,991 (90.4%) | 23,376 (96.1%) | 18,511 (91.3%) | 19,593 (96.7%) |
| **otoArg** | 23,623 | 21,339 (90.3%) | 22,699 (96.1%) | 18,475 (91.2%) | 19,576 (96.6%) |
| **copChr** | 22,977 | 20,862 (90.8%) | 22,062 (96.0%) | 18,440 (91.0%) | 19,505 (96.2%) |
| **rhaChi** | 24,410 | 21,957 (90.0%) | 23,375 (95.8%) | 18,410 (90.8%) | 19,531 (96.4%) |
| **rhaChi2** | 23,075 | 20,817 (90.2%) | 22,104 (95.8%) | 18,404 (90.8%) | 19,496 (96.2%) |

**Supplemental Table S4.** Estimated gene sequence conservation by ODGI's presence-absence variation analysis. PAV analysis was restricted to gene sets meeting criteria for sufficient sequencing coverage, avoidance of complex graph regions and removal of falsely annotated TEs. A different set of reliable genes was used for each sample (first three columns), as well as a universally reliable set for all samples (last two columns). The table counts the number of genes showing 90% sequence conservation or above. Backbone assemblies are shaded in gray.

| | Observed (O) | Genomewide (G) | Expected (E) | O/G ratio | O/E ratio |
|---|---|---|---|---|---|
| **All Transposons (TEs)** | 74.65 | 36.55 | 42.46 | 2.04 | 1.76 |
| **All TEs w/o Unknown** | 62.25 | 28.30 | 34.57 | 2.20 | 1.80 |
| **DNA** | 19.55 | 12.09 | 12.08 | 1.62 | 1.62 |
| **LINE** | 16.82 | 8.32 | 10.05 | 2.02 | 1.67 |
| **LTR** | 25.31 | 7.38 | 12.18 | 3.43 | 2.08 |
| **SINE** | 0.35 | 0.40 | 0.30 | 0.88 | 1.17 |
| **Retroposon** | 0.50 | 0.24 | 0.33 | 2.08 | 1.52 |
| **Helitron** | 0.20 | 0.06 | 0.10 | 3.33 | 2.00 |
| **Unknown** | 12.92 | 8.54 | 8.50 | 1.51 | 1.52 |

**Supplemental Table S5.** Fold enrichment of TEs in SV regions of *Astatotilapia calliptera*. The percentage sequence of non-overlapping TE regions are shown for the structural variants (O) and whole genome (G). The E column are mean values estimated from 100 randomly shuffled sets of SV coordinates that still maintained proximity to the SV regions, in order to create more realistic baselines than the genomewide values.

(a) Genomewide

| | astCal | mayZeb | troMau | aulStu | otoArg | copChr | rhaChi | rhaChi2 |
|---|---|---|---|---|---|---|---|---|
| **All Transposons (TEs)** | 36.55 | 38.29 | 37.54 | 37.13 | 37.09 | 36.32 | 37.49 | 36.96 |
| **All TEs w/o Unknown** | 28.30 | 29.36 | 28.71 | 28.20 | 28.39 | 28.03 | 29.02 | 28.65 |
| **DNA** | 12.09 | 12.23 | 12.08 | 12.12 | 12.03 | 11.81 | 12.22 | 12.12 |
| **LINE** | 8.32 | 8.77 | 8.50 | 8.64 | 8.46 | 8.47 | 8.60 | 8.46 |
| **LTR** | 7.38 | 7.65 | 7.48 | 7.53 | 7.32 | 7.21 | 7.57 | 7.45 |
| **SINE** | 0.40 | 0.40 | 0.40 | 0.40 | 0.39 | 0.39 | 0.40 | 0.39 |
| **Retroposon** | 0.24 | 0.35 | 0.26 | 0.24 | 0.29 | 0.27 | 0.28 | 0.28 |
| **Helitron** | 0.06 | 0.18 | 0.21 | 0.10 | 0.12 | 0.09 | 0.18 | 0.17 |
| **Unknown** | 8.54 | 9.22 | 9.11 | 8.61 | 8.98 | 8.57 | 8.76 | 8.61 |

(b) Structural variants

| | astCal | mayZeb | troMau | aulStu | otoArg | copChr | rhaChi | rhaChi2 |
|---|---|---|---|---|---|---|---|---|
| **All Transposons (TEs)** | 74.65 | 73.29 | 72.26 | 73.85 | 74.25 | 75.74 | 74.82 | 76.60 |
| **All TEs w/o Unknown** | 62.25 | 61.74 | 59.99 | 62.78 | 62.79 | 65.10 | 62.79 | 65.32 |
| **DNA** | 19.55 | 19.32 | 18.36 | 19.36 | 19.11 | 19.59 | 19.76 | 20.39 |
| **LINE** | 16.82 | 17.85 | 16.91 | 18.21 | 18.40 | 19.34 | 17.75 | 18.36 |
| **LTR** | 25.31 | 24.09 | 23.82 | 24.88 | 24.92 | 25.81 | 24.77 | 26.19 |
| **SINE** | 0.35 | 0.33 | 0.32 | 0.31 | 0.31 | 0.32 | 0.29 | 0.29 |
| **Retroposon** | 0.50 | 0.47 | 0.41 | 0.41 | 0.41 | 0.44 | 0.43 | 0.43 |
| **Helitron** | 0.20 | 0.20 | 0.68 | 0.17 | 0.20 | 0.16 | 0.35 | 0.22 |
| **Unknown** | 12.92 | 12.02 | 12.74 | 11.56 | 11.95 | 11.15 | 12.59 | 11.84 |

**Supplemental Table S6.** Genomewide and structural variant TE composition across Lake Malawi cichlid assemblies.

### Supplemental Methods

#### Long read genome assemblies

The six new long read assemblies in this study were generated from DNA extracted from aquaria grown fish specimens. The *Tropheops* sp. “mauve” and two *Rhamphochromis* sp. "Chilingali" individuals (one male, one female) were reared by members of the Santos Lab at the local fish facility in the University of Cambridge. *Otopharynx argyrosoma* and *Copadichromis chrysonotus* were provided by Dr. Hannes Svardal’s lab at the University of Antwerp in Belgium, while *Aulonocara stuartgranti* were sourced from Professor George Turner at Bangor University, Wales. Pacific Biosciences (PacBio) sequencing was used for *Tropheops* sp. “mauve” (troMau), *Aulonocara stuartgranti* (aulStu) and the male *Rhamphochromis* sp. “Chilingali” (rhaChi), while Oxford Nanopore (ONT) was used for *Copadichromis chrysonotus* (copChr), *Otopharynx argyrosoma* (otoArg) and the female *Rhamphochromis* (rhaChi2). Depending on tissue availability, frozen tissue from either fin, muscle or gill tissue, with the variations in DNA extraction and library preparation protocols described below.

##### Pacific Biosciences

**Genomic DNA extraction**

Tissue samples were collected from muscle for *Tropheops* sp. “mauve” (troMau) and the male *Rhamphochromis* sp. “Chilingali” (rhaChi) sample, while fin and tail clips were used for *Aulonocara stuartgranti* (aulStu). High molecular weight DNA (HMW DNA) was extracted using the Bionano Genomics: IrysPrep® Animal Tissue DNA Isolation Soft Tissue protocol, which involved tissue disruption, cell lysis and DNA purification steps. The quantity of extracted genomic DNA was evaluated with the HS Qubit DNA kit and QC-assessed with the Femto Pulse instrument (Agilent).

**Library preparation**

Pacific Biosciences (PacBio) SMRT sequencing was performed with CLR (Continuous Long Reads) technology for the HWM DNA, following the official SMRTbell® Express Template Preparation Kit 2.0 protocol, with specific modifications to the DNA shearing step and library size selection using a Bluepippin instrument (Sage Science) depending on the sample.

- troMau: DNA shearing was not performed, because the sample was slightly degraded. Library size of 20 kb selected.
- aulStu: DNA sheared using a 26 gauge needle (5 passes through the needle). Library size of 7 kb selected
- rhaChi: DNA sheared using Megaruptor 3 at speed setting 5. Library size of 20 kb.

The prepared DNA libraries were sequenced on the Sequel II instrument using Sequencing kit v0.9 / Binding Kit v0.9. Contigs were generated from the reads using FALCON and FALCON-Unzip assembler software.

##### Oxford Nanopore

**DNA extraction**

*Otopharynx argyrosoma*, otoAtg: Fin tissue was pulverised in a mortar with liquid nitrogen, after which DNA was extracted using the QIAGEN Genomic Tip 100/G Kit (10243) with minor modifications to manufacturer instructions. After elution from the Genomic Tip column, the sample was divided into 4 x 1.25 mL aliquots in 2.0 mL DNA LoBind® tubes using wide-bore tips and the DNA precipitated with isopropanol, washed twice with 70% ethanol and each aliquot resuspended in 50uL elution buffer.

*Copadichromis chrysonotus*, copChr: Fin tissue was pulverised in a mortar with liquid nitrogen. Released cells were lysed in 5 mL Lysis Buffer (100 mM NaCl, 10 mM Tris pH, 25 mM EDTA pH, 0.5% SDS) with 2 uL RNaseA (100 mg/ml) at 37°C for 1 h with gentle inversion mixing every 20 minutes, followed by 50 uL Proteinase K (20 mg/ml) digestion at 55°C for 2 hours with 10 rpm in a rotating incubator. DNA was then extracted using Phenol-Chloroform-Isoamyl alcohol pH 8 in combination with MaXtract High Density 15 ml tubes (Qiagen) according to manufacturer instructions. The DNA was then precipitated with isopropanol in the presence of NaCl, spooled and transferred to DNA LoBind® tubes, washed twice with 70% ethanol, air dried for 10 minutes and resuspended in a 100 uL elution buffer. DNA was size selected on Bluepippin at >15kb followed by magnetic bead purification.

*Rhamphochromis* female, rhaChi2: Gill tissue was cut into smaller pieces and digested for 2.5h in ALT Buffer with 20 uL Proteinase K (20 mg/ml) at 56°C. Undigested cell debris removed by a very gentle centrifugation and supernatant transferred with wide-bore tip to DNA LoBind® tube. 4 uL RNaseA (100 mg/ml) was added and incubated for 10 minutes at room temperature. Tween 20 (to a 0.1% final concentration) was added with an equal volume of resuspended SPRIselect beads and incubated on a tube rotator at 10 rpm for 20 minutes. The beads were captured on a magnet and washed twice with fresh 70% ethanol for 30 seconds, briefly air dried, then removed from the magnet and resuspended with 55 uL elution buffer and mixed by gently tapping on the tube and incubated for a few minutes at room temperature. DNA was then separated from the beads on the magnet and supernatant transferred using wide bore tips to the new tube. Elution was repeated but collected into separate tubes. DNA was size selected on Bluepippin at >20kb followed by magnetic bead purification.

**Library preparation and genome assembly**

Prior to library preparation, DNA was QC assessed using Qubit (dsDNA BR Assay Kit), Nanodrop and Tapestation (Genomic DNA ScreenTape). Sequencing libraries were prepared from 1-5 ug of material using Ligation Sequencing Kit (SQL-LSK109) and sequenced on R9 MinION flow cells with minor modifications. End repair was performed at 20°C for 30 minutes, followed by 30 minutes at 65°C, after which adapter ligation was performed for 1 hour. Guppy v5.0.11 was used to call bases, and the resulting read sets were assembled into contigs with Shasta v0.7.0 (commit b64c4ad0755a6e0d5ba5ef9c1cffead5dc6fadc0) [182] under default settings.

#### PCR experiments

**Fish maintenance and euthanasia**

*Astatotilapia calliptera*, *Tropheops sp. "mauve"* and *Rhamphochromis sp. "chilingali"* animals were grown in 220 Litre tanks, with pH 8, at approximately 28°C, and with a 12h dark/light cycle. Males and females of each species were housed only with conspecifics. Feeding, housing, and handling were conducted in strict adherence to local regulations. Fish were fed twice a day with cichlid flakes and pellets (Vitalis). Tank environment was enriched with plastic plants, plastic hiding tubes, and sand substrate.

Aquaria grown animals were euthanized with 1 g/L MS-222 (Ethyl 3-aminobenzoate methanesulfonate, Merck #E10521) and subsequent exsanguination by cutting the gill arches, in accordance with local regulations. Afterwards, required tissues were carefully dissected, swiftly snap frozen in dry ice and stored at approximately -80°C.

**Lysis and DNA extraction**

DNA was extracted from frozen fins or muscle tissue using the QIAamp DNA Mini kit (Qiagen, #51304), according to manufacturer's instructions. A small portion of fin or muscle tissue was lysed using the lysis buffer supplied in the kit, supplemented with Proteinase K. Lysis was performed at 37°C for 1-2 h. The quality and purity of the extracted DNA was checked using a Nanodrop 2000 (Thermo Fischer Scientific) and on an 1% agarose gel. 1-4 ng of genomic DNA was subsequently used as a template in each PCR reaction. DNA was extracted from two distinct sets of animals: 1) from the same fin tissue of the wild-caught animals whose DNA was sequenced to generate the genome assemblies (one male each of *Tropheops sp. "mauve"*, *Otopharynx argyrosoma*, and *Copadichromis chrysonotus*, and one male and one female of *Rhamphochromis sp. "chilingali"*, no fin tissue leftover for the other species); and 2) from the fins or muscle of aquaria grown individuals (five males and five females of *Astatotilapia calliptera salima*, nine *Tropheops sp. "mauve"* males, and five females and one male of *Rhamphochromis sp. "chilingali"*).

**Experimental validation of SVs by PCR and Sanger sequencing**

PCRs to validate SVs were performed using Taq DNA Polymerase (NEB #M0267) following manufacturer's instructions. 10 µl reactions were prepared and 1-4 ng of genomic DNA were used as template. Primers used to amplify distinct SVs in the vicinity of protein-coding genes are listed in below. Taq PCR reactions were performed as follows: 95°C for 1 minute; 35 cycles of 95°C for 30 seconds, 60°C for 30 seconds, and 68°C for varying periods of time; and a final cycle of 68°C for 5 minutes. PCR products were run on a 1% agarose gel stained with SYBR Safe (Thermo Fischer Scientific, #S33102) at 120V for 50 minutes.

For subsequent validation of amplicons by Sanger sequencing, PCRs were repeated using high-fidelity Q5 DNA Polymerase (NEB, #M0491), according to manufacturer's instructions. 50 µL reactions were prepared and 6-8 ng of genomic DNA was used as template. Primers used are the same used for Taq DNA Polymerase PCRs, and are listed below. When sequencing was inefficient, additional primers were designed, which include M13 primer sites for more efficient sequencing. 5 µl of the PCR product were run on a 1% agarose gel stained with SYBR Safe to confirm the presence of bands of expected size, and the remaining PCR product was sent to Azenta Life Sciences for Sanger sequencing with the primers listed or M13 universal primers. For animals heterozygous for a particular SV, as observed by the presence of two bands on an agarose gel, the entire PCR product was run on a 1% agarose gel and both bands were extracted using the QIAquick Gel Extraction kit (Qiagen, #28704), and each sent to Azenta Life Sciences for Sanger sequencing.

### Table: PCR primers for selected bubbles

| Oligo ID | Fw/Rev | Sequence (5' to 3') | SV ID (proximal protein-coding gene) | PCR Extension Time | Expected PCR Product Size (in base pairs)* astCal | troMau | otoArg | copChr | rhaChi | rhaChi2 | Notes |
|---|---|---|---|---|---|---|---|---|---|---|---|
| MVA380 | Fw | CAGTGGAGGAGGATCTCAGC | s180017 (*ENSACLG00000015768* aka *nsd1b*) | 1 min 30 sec | 944 | 729 | 944 | 944 | 729 | 729 | |
| MVA381 | Rev | TTTTGCTTTTCTGCCTCGAT | | | | | | | | | |
| MVA391 | Fw | TGCAGAAGACGCTCTGATCT | s438346/s54791 (ENSACLG00000021983) | 1 min 30 sec | 1306 | 169 | 169 | 1306 | 1306 | 1306 | |
| MVA392 | Rev | TGCAGCAACATTTCAAAACAA | | | | | | | | | |
| MVA395 | Fw | GAGACTCTCAGCCGTTTTACG | s461001 (*ENSACLG00000002767* aka *chchd3b*) | 1 min | 281 | 588 | 588 | 588 | 281 | 281 | |
| MVA396 | Rev | TCCACACACTGCATGTAGGC | | | | | | | | | |
| MVA399 | Fw | AAATGGGTCATTTGGAGCTG | s175418 (*ENSACLG00000017346* aka *kcnab3*) | 40 sec | 446 | 133 | 446 | 446 | 133 | 133 | |
| MVA400 | Rev | CCCAATAGTGTCCACTGACATC | | | | | | | | | |
| MVA401 | Fw | CTGAACCCGTTTCTCACACA | s120363 (*ENSACLG00000027791* aka *ptges*) | 40 sec | 460 | 143 | 143 | 143 | 460 | 460 | |
| MVA402 | Rev | CCAGTGCTGGGTTTTGAGAT | | | | | | | | | |
| MVA409 | Fw | TCAGGTAGAGGGCAGGTGTT | s165551 (*ENSACLG00000013807* aka *gabrr3b*) | 1 min | 852 | 338 | 338 | 338 | 852 | 852 | |
| MVA410 | Rev | CTGTCACGTCATTGGCTGAT | | | | | | | | | |
| MVA413 | Fw | TGAGGAGGAAGAGGATTTGG | s473871 (*ENSACLG00000014403* aka *adam17b*) | 40 sec | 207 | 413 | 207 | 207 | 413 | 413 | |
| MVA414 | Rev | CTGCAGTCAGCTGGGTTTTT | | | | | | | | | |
| MVA415 | Fw | CATGCTTTCTGCATGCATCT | s171149 (*ENSACLG00000026652* aka *dgat2*) | 40 sec | 558 | 327 | 558 | 558 | 327 | 327 | |
| MVA416 | Rev | ACGTGCTTGGCTTTCAAGAG | | | | | | | | | |
| MVA421 | Fw | GGAGGGAAAACAGCCAAAAT | s88741 (*ENSACLG00000022427* aka *mitfa*) | 1 min 35 sec | 1037 | 555 | 1037 | 1037 | 1037 | 1037 | |
| MVA422 | Rev | CACCGACGTAATGGGAAACT | | | | | | | | | |
| MVA431 | Fw | AGGTTCCTGCTGAAGGTCAA | s483914 (*ENSACLG00000025203* aka *mfsd4aa*) | 1 min | 230 | 592 | 230 | 230 | 230 | 230 | |
| MVA432 | Rev | GGTCGACGGAATTCTCATGT | | | | | | | | | |
| MVA435 | Fw | GGATTTGGCACTACCTTCCA | s336440 (*ENSACLG00000006581* aka *klhl7*) | 1 min 35 sec | 234 | 1519 | 234 | 234 | 234 | 234 | |
| MVA436 | Rev | ATGCCCCAACAACTTGAAAA | | | | | | | | | |
| MVA439 | Fw | GCCACCACATGTCTCTCAAA | s457179 (*ENSACLG00000020186* aka *rpgrip1*) | 1 min 35 sec | 215 | 1234 | 215 | 215 | 215 | 215 | |
| MVA440 | Rev | CCACAGACTGCTTGACCTGA | | | | | | | | | |
| MVA445 | Fw | ATGTCCATCTTGAGGGCTGT | s473698 (*ENSACLG00000017196* aka *anpepb*) | 1 min 10 sec | 218 | 863 | 218 | 218 | 218 | 218 | |
| MVA446 | Rev | GCATCAAGACGTACCGTCAA | | | | | | | | | |
| MVA449 | Fw | GCGCATCTGGACTCATTTGT | s256152 (*ENSACLG00000027349* aka *dtnba*) | 1 min 10 sec | 826 | 193 | 826 | 193 | 193 | 193 | |
| MVA450 | Rev | TGTTTTTCAGCCAACTGTGG | | | | | | | | | |
| MVA451 | Fw | ACCCAAAGGAGACGAAGAGG | s423054 (*ENSACLG00000015018* aka *lcp1*) | 1 min 10 sec | 640 | 1062 | 640 | 1062 | 1062 | 1062 | |
| MVA452 | Rev | CTCGCTTCAGGTCGTCTTTC | | | | | | | | | |
| MVA457 | Fw | GTTCTCAGCGTTTGGCTGAT | s364746 (*ENSACLG00000024576* aka *tprg1*) | 1 min 10 sec | 598 | 289 | 598 | 598 | 598 | 598 | |
| MVA458 | Rev | ACAACAGCCCCAGCTTCTC | | | | | | | | | |
| MVA459 | Fw | **GTAAAACGACGGCCAG**GGCGTGTACTCTGAACCCGTTTCTCACACA | s120363 (*ENSACLG00000027791* aka *ptges*) | 1 min | 518 | 201 | 201 | 201 | 518 | 518 | With M13 Fw (Bold) |
| MVA460 | Rev | **CAGGAAACAGCTATGAC**AGGTCTCTGATAACACCAGTGCTGGGTTTTGAGAT | | | | | | | | | With M13 Rev (Bold) |
| MVA461 | Fw | **GTAAAACGACGGCCAG**GAGACTCTCAGCCGTTTTACG | s461001 (*ENSACLG00000002767* aka *chchd3b*) | 1 min | 314 | 621 | 621 | 621 | 314 | 314 | With M13 Fw (Bold) |
| MVA462 | Rev | **CAGGAAACAGCTATGAC**TCCACACACTGCATGTAGGC | | | | | | | | | With M13 Rev (Bold) |
| MVA463 | Fw | **GTAAAACGACGGCCAG**GCGCATCTGGACTCATTTGT | s256152 (*ENSACLG00000027349* aka *dtnba*) | 1 min | 879 | 246 | 879 | 246 | 246 | 246 | With M13 Fw (Bold) |
| MVA464 | Rev | **CAGGAAACAGCTATGAC**AATTAAATATTTTTGCTCAATGTTTTTCAGCCAACTGTGG | | | | | | | | | With M13 Rev (Bold) |
| MVA465 | Fw | **GTAAAACGACGGCCAG**AGGTTCCTGCTGAAGGTCAA | s483914 (*ENSACLG00000025203* aka *mfsd4aa*) | 1 min | 263 | 625 | 263 | 263 | 263 | 263 | With M13 Fw (Bold) |
| MVA466 | Rev | **CAGGAAACAGCTATGAC**GGTCGACGGAATTCTCATGT | | | | | | | | | With M13 Rev (Bold) |

*PCR product size indicated for each of the species used for PCR. Species notation used is the same as in the main text.
